# CLN6 is a glycerophosphoglycerol epimerase in bis(monoacylglycero)phosphate biosynthesis

**DOI:** 10.64898/2026.10.07.757350

**Authors:** Pradeep K. Sheokand, Andrew M. James, Christian Posern, Amit Singh Sahrawat, Johannes Breithofer, Joe Ganellin, McKayla K. Proulx, Susannah K. Leese, Keira Turner, Denis Lacabanne, Dominik Bulfon, Benjamin Jenkins, Joris van der Kleij, Harvey E. Johnston, Samuel O. Lord, Maya K. Basran, Chak Shun Yu, Clarissa D. Booth, Jill M. Weimer, Albert Koulman, Marja Jäättelä, Edmund R. S. Kunji, David N. Palmer, Nadia L. Mitchell, Julien Prudent, Robert Zimmermann, Karl Gruber, Angela Schulz, Michael P. Murphy, Kasparas Petkevicius

## Abstract

Lysosomal catabolism depends on bis(monoacylglycero)phosphate (BMP), a phospholipid with atypical *S,S* stereochemistry of unknown biosynthetic origin. CLN8 and CLN5 catalyze sequential acylation steps in BMP synthesis, and their loss of function causes Batten disease, a group of inherited neurodegenerative lysosomal storage disorders. Here we identify CLN6, a biochemically uncharacterized endoplasmic reticulum protein whose loss also causes Batten disease, as the calcium-dependent glycerophosphoglycerol epimerase that establishes BMP stereochemistry upstream of CLN8. CLN6 deficiency depletes BMP in cells, mice, and patient lysosomes, and reduces lysosomal hydrolase levels. BMP-precursor supplementation bypasses CLN6, restores BMP and increases hydrolase abundance in CLN6-deficient cells and mice. These findings place three Batten disease proteins in a single biosynthetic pathway and reveal BMP deficiency as a reversible cause of disrupted lysosomal hydrolase homeostasis.

**One sentence summary:** CLN6 establishes the *S,S* stereochemistry of bis(monoacylglycero)phosphate and links its synthesis to lysosomal hydrolase homeostasis.

## Main text

Batten disease is a group of inherited neurodegenerative disorders characterized by the accumulation of undegraded material in lysosomes (*1*). Several forms result from deficiencies of soluble lysosomal hydrolases, including palmitoyl-protein thioesterase 1 (PPT1/CLN1), tripeptidyl peptidase 1 (TPP1/CLN2) and cathepsin D (CTSD/CLN10), whereas others arise from defects in membrane proteins of unknown biochemical function (*1*). One such protein is CLN6, a multipass endoplasmic reticulum (ER) membrane protein with no human paralog or defined protein family (*2–4*). Loss of CLN6 depletes PPT1, TPP1, CTSD and other hydrolases from lysosomes without reducing their transcript levels, while lysosomal membrane proteins are preserved (*5–7*). This selective depletion has been attributed to a putative role for a CLN6-CLN8 complex in transporting newly synthesized lysosomal hydrolases from the ER to the Golgi (*5*).

Depletion of bis(monoacylglycero)phosphate (BMP), a phospholipid of lysosomal intralumenal membranes that promotes lipid catabolism, is another feature of several forms of Batten disease (*8–13*). BMP is stereochemically unusual: whereas other mammalian glycerophospholipids carry an *R*-configured glycerol backbone, BMP contains two *S*-configured glycerol moieties (*S,S*-BMP) (*14–16*). This configuration confers resistance to lysosomal phospholipases, allowing BMP to persist in a degradative compartment (*17*, *18*). Because its precursor, phosphatidylglycerol, has an *R,S* configuration (*R,S*-PG), BMP synthesis requires stereoinversion of one glycerol moiety from *R* to *S* (*19*). The mechanism and enzyme responsible for this stereoinversion have remained a central unresolved problem in lipid biochemistry for over five decades, since BMP’s unusual stereochemistry was first established (*14*, *15*).

We and others recently identified two Batten disease proteins that catalyze successive acyl transfer reactions in BMP synthesis. The ER acyltransferase CLN8 uses acyl-CoA to convert *S,S*-glycerophosphoglycerol (*S,S*-GPG) to *S,S*-lysophosphatidylglycerol (*S,S*-LPG), which the lysosomal transacylase CLN5 then converts to BMP (*8*, *12*, *13*). Both enzymes act on substrates that already carry the *S,S* configuration, placing the missing stereoinversion step upstream of CLN8 (Fig. 1A).

**Fig. 1.**
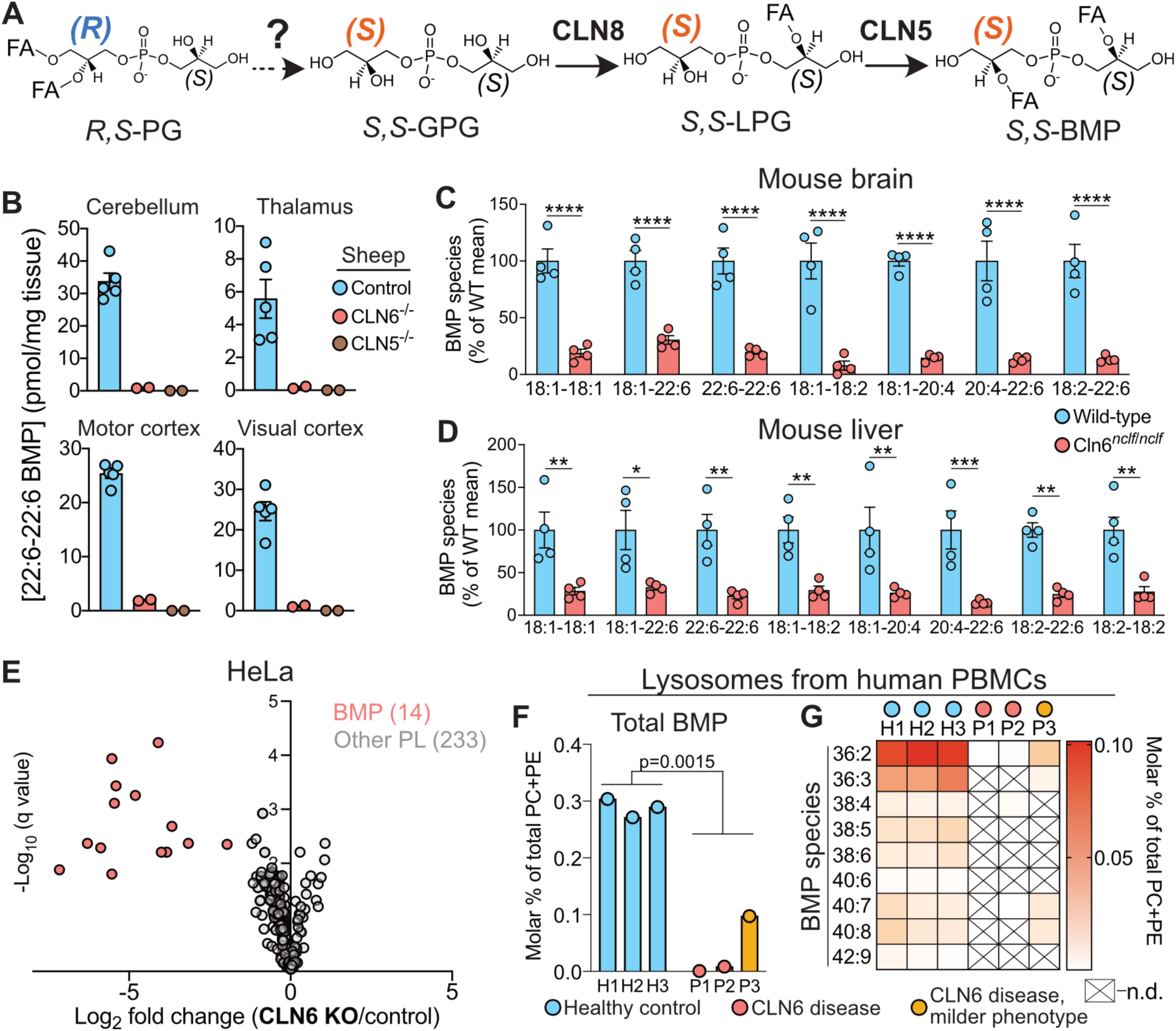
CLN6 deficiency depletes BMP in animal models, cultured cells and patient lysosomes. (**A**) Schematic of BMP biosynthesis, showing the unresolved formation of *S,S*-glycerophosphoglycerol (*S,S*-GPG) from *R,S*-phosphatidylglycerol (*R,S*-PG), followed by sequential acylation by CLN8 and CLN5. LPG, lysophosphatidylglycerol; FA, fatty acyl chain. (**B**) Abundance of 22:6–22:6 BMP in the indicated brain regions from 12-month-old female control (n = 5), CLN6-deficient (n = 2) and CLN5-deficient (n = 2) sheep, measured by targeted LC-MS/MS and expressed per milligram of tissue. (**C** and **D**) Abundance of the indicated BMP species in brain (**C**) and liver (**D**) from 6-month-old female wild-type and *Cln6^nclf/nclf^* mice, measured by targeted LC-MS/MS and expressed as a percentage of the wild-type mean for each species (n = 4 mice per group). (**E**) Volcano plot of phospholipid abundance in CLN6 knockout (KO) relative to control HeLa cells, measured by untargeted LC-MS (n = 3 biological replicates per group). Each point represents a lipid species; 14 BMP species are highlighted in red, and 233 other phospholipid species are shown in grey. (**F** and **G**) Total BMP abundance (**F**) and abundance of individual BMP species (**G**) in lysosomes isolated from peripheral blood mononuclear cells of three healthy young adult controls (H1–H3) and three children with CLN6 Batten disease (P1–P3). P3 had a milder disease phenotype. Lipids were measured by untargeted LC-MS, and BMP abundance is expressed as a molar percentage of the combined phosphatidylcholine (PC) and phosphatidylethanolamine (PE) abundance. In (**G**), crossed cells indicate species not detected (n.d.). Data in (**B**–**D**) are means ± SEM, with points representing individual animals. In (**C**) and (**D**), *P < 0.05, **P < 0.01, ***P < 0.001 and ****P < 0.0001 by two-way ANOVA with Šídák’s multiple-comparisons test. In (**E**), P values from multiple t tests were adjusted using the two-stage Benjamini–Krieger–Yekutieli false-discovery rate procedure. In (**F**), the indicated P value compares controls with patients using an unpaired two-tailed Student’s t test.

Three observations suggested a role for CLN6 in BMP biosynthesis. First, CLN6 loss produces molecular phenotypes closely resembling those of CLN8 deficiency, and combined loss of both proteins does not further accelerate neurodegeneration in mice, consistent with their action in a shared pathway (*5*). Second, sheep carrying loss-of-function mutations in CLN5 or CLN6 develop nearly identical brain pathology and disease progression (*20*, *21*); CLN5 deficiency causes near-complete BMP loss in sheep tissues (*22*). Third, CLN6 homologues are absent from yeast and flies, neither of which synthesizes BMP (*23*). Here we identify CLN6 as the glycerophosphoglycerol epimerase that establishes the unusual *S,S* stereochemistry of BMP.

## Results

### CLN6 deficiency causes BMP depletion

To investigate the role of CLN6 in BMP synthesis, we first measured BMP and its precursor LPG in multiple brain regions from control, CLN5-deficient and CLN6-deficient sheep (*20*). As previously reported, CLN5-deficient sheep showed loss of BMP across all regions analyzed, accompanied by elevated LPG levels, consistent with impaired conversion of LPG to BMP (Fig. 1B, fig. S1, A and B) (*22*). CLN6-deficient sheep likewise showed marked BMP depletion and LPG accumulation in every region examined, but retained a small residual BMP pool that was absent in CLN5-deficient sheep (Fig. 1B, fig. S1, A and B).

We next analyzed BMP and LPG in brains and livers from homozygous *Cln6^nclf/nclf^* mice, which carry a frameshift mutation in *Cln6* and are widely used as a model of CLN6 Batten disease (*24*, *25*). Relative to WT controls, all measured BMP species were strongly reduced in both tissues, whereas LPG species were elevated (Fig. 1, C and D, fig. S1C). Brain GPG was also increased in *Cln6^nclf/nclf^* mice compared to controls, paralleling our previous observations in CLN8-deficient mice (fig. S1D) (*8*). The effects on LPG, however, were opposite in the two models: LPG was elevated in *Cln6^nclf/nclf^* brain but strongly reduced in CLN8-deficient brain (*8*, *13*), suggesting that CLN6 and CLN8 act at distinct steps of BMP biosynthesis (fig. S1C). Notably, residual BMP species bearing different acyl chains in CLN6-deficient sheep and mouse tissues showed altered relative product-ion abundances by LC-MS/MS compared with the corresponding species in controls, suggesting a structural difference (fig. S2, A to C).

To test whether BMP depletion is a cell-autonomous consequence of CLN6 loss, we generated a polyclonal CLN6-knockout (CLN6 KO) HeLa population by CRISPR and compared its phospholipidome with that of an isogenic wild-type control population (fig. S3, A to C). Consistent with the sheep and mouse data, all measured BMP species were strongly decreased in CLN6 KO cells, whereas other major membrane phospholipids were relatively unchanged (Fig. 1E, fig. S3D).

Finally, we used a recently described tagless Lyso-IP method to isolate lysosomes from peripheral blood mononuclear cells (PBMCs) of three children with CLN6 Batten disease and three healthy young adult controls (*26*). Two patients had a typical late infantile presentation, and one (P3) had an atypical, milder form. Untargeted lipidomics of the isolated lysosomes, with BMP normalized to the major membrane phospholipids phosphatidylcholine (PC) and phosphatidylethanolamine (PE) (fig. S2D), showed that BMP was almost entirely absent from the lysosomes of the two patients with typical disease and strongly reduced in P3 relative to controls (Fig. 1F). Together, these findings establish CLN6 as a conserved determinant of BMP abundance across cultured cells, animal disease models and patient material.

### CLN6 catalyzes GPG stereoinversion

To determine where CLN6 acts in BMP biosynthesis, we used stable isotope tracing in cultured cells. We previously used an *R,rac*-PG tracer containing a deuterium-labeled racemic (*rac*; 50% *R*-, 50% *S*-) glycerol headgroup and a C16:0/C18:1 *R*-glycerol backbone (d_5_-PG) to define the CLN8 reaction (*8*). In that setting, loss of CLN5 blocked the final step of BMP synthesis and trapped the upstream CLN8 product, LPG, in which the deuterium label was distributed symmetrically between the headgroup and backbone — demonstrating that CLN8 acylates a symmetrical *S,S*-GPG substrate (Fig. 2A) (*8*).

**Fig. 2.**
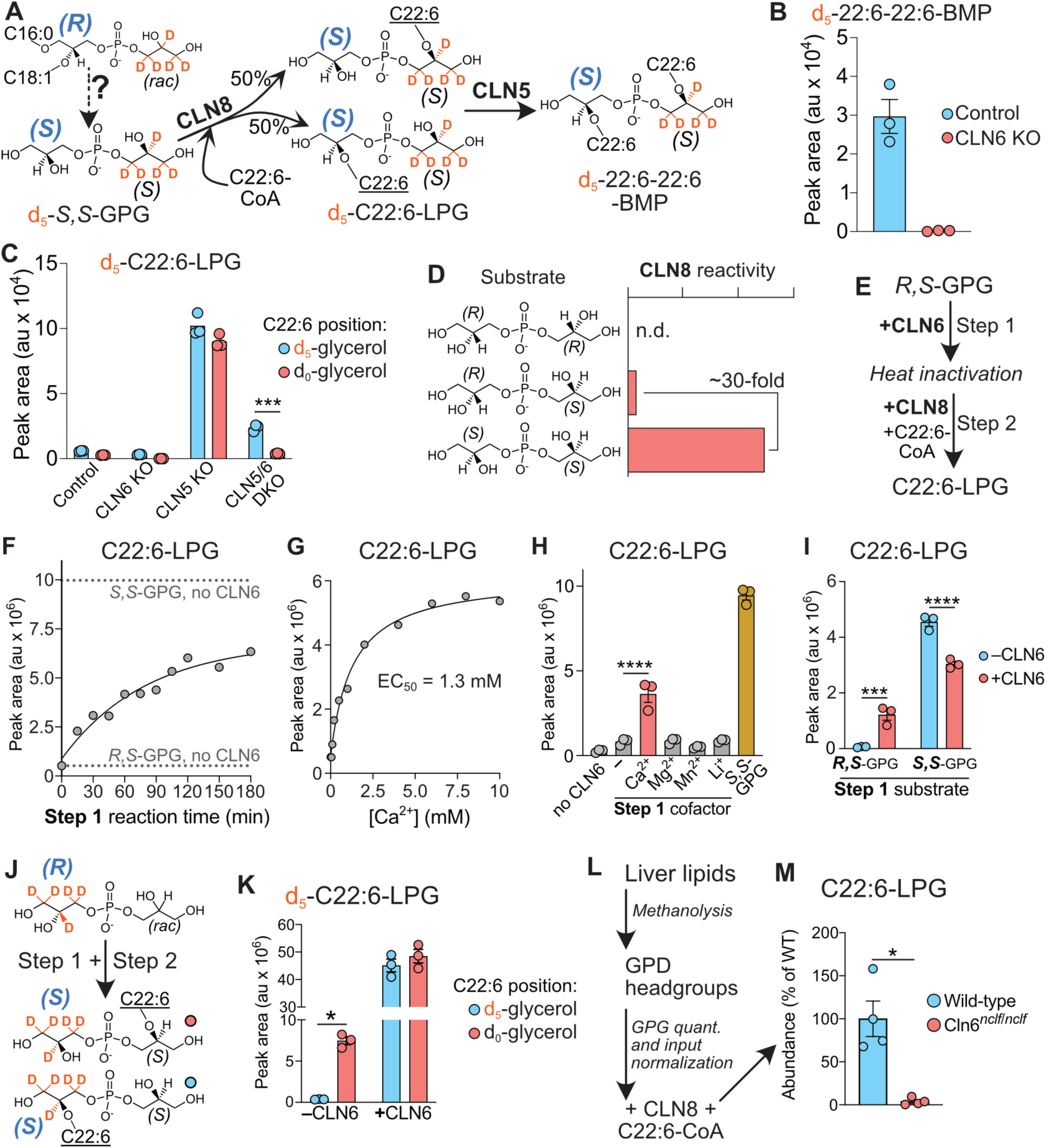
CLN6 is a Ca^2+^-dependent *R,S*-GPG epimerase. (**A**) Schematic tracing the conversion of d_5_-16:0/18:1-PG to d_5_-22:6–22:6 BMP in cells. CLN8 acylation of d_5_-*S,S*-GPG produces LPG with the acyl chain on either the labeled or unlabeled glycerol moiety. (**B**) d_5_-22:6–22:6 BMP abundance in control and CLN6 KO HeLa cells following a 4-hour pulse with d_5_-PG (n = 3). (**C**) Abundance of d_5_-C22:6-LPG carrying C22:6 on the d_5_- or d_0_-glycerol moiety in control, CLN6 KO, CLN5 KO and CLN5/6 double-knockout (DKO) HeLa cells following a 4-hour pulse with d_5_-PG (n = 3). (**D**) Schematic comparison of CLN8 reactivity towards GPG stereoisomers. n.d., not detected. (**E**) Two-step assay coupling CLN6-mediated stereoconversion of *R,S*-GPG to CLN8-dependent acylation. CLN6 is heat-inactivated after step 1, and C22:6-LPG formation in step 2 provides the reaction readout. (**F**) C22:6-LPG formation following the indicated durations of step 1. The curve shows a one-phase association fit (R^2^ = 0.96). Upper and lower dotted lines indicate CLN8-only reactions supplied with *S,S*-GPG or *R,S*-GPG, respectively. (**G**) Ca^2+^ concentration dependence of the coupled assay, with Ca^2+^ varied during step 1 (apparent EC_50_ = 1.3 mM). (**H**) C22:6-LPG formation with the indicated metal ions supplied at 5 mM during step 1; no-CLN6 and *S,S*-GPG controls are also shown. (**I**) C22:6-LPG formation from *R,S*-GPG or *S,S*-GPG incubated with or without CLN6 in step 1. (**J**) Schematic of stereoconversion and subsequent acylation of d_5_-*R,rac*-GPG, labeled on the glycerol moiety opposite to that labeled in (**A**). (**K**) Abundance of d_5_-C22:6-LPG carrying C22:6 on the d_5_- or d_0_-glycerol moiety after reactions with or without CLN6 in step 1. (**L**) Workflow for assessing the stereochemistry of GPG released from liver lipids by methanolysis. GPG was quantified and equal amounts were supplied to CLN8 with C22:6-CoA. (**M**) C22:6-LPG formation from liver-derived GPG from 6-month-old female wild-type and *Cln6^nclf/nclf^* mice, expressed as a percentage of the wild-type mean (n = 4 mice per group). Data are means ± SEM where error bars are shown. Points represent biological replicates in (**B**, **C**), individual enzymatic reactions in (**F**–**I**, **K**) and individual mice in (**M**). In (**C**, **I**, **K**), *P < 0.05, ***P < 0.001 and ****P < 0.0001 by two-way ANOVA with Šídák’s multiple-comparisons test. In (**H**), ****P < 0.0001 by one-way ANOVA with Tukey’s multiple-comparisons test. In (**M**), *P < 0.05 by an unpaired two-tailed Student’s t test. Purified CLN6 assays are representative of experiments using three independent protein purifications.

We applied the same strategy to CLN6-deficient cells. Using CRISPR, we generated parallel polyclonal CLN6 KO, CLN5 KO and CLN5/6 double KO HeLa cell populations and traced the metabolism of d_5_-PG (fig. S3, E to G). CLN6 KO cells failed to convert d_5_-PG into d_5_-22:6-22:6 BMP, indicating that CLN6 is required for efficient BMP synthesis under these conditions (Fig. 2B). We then used CLN5 deficiency to trap d_5_-C22:6-LPG, whose formation requires complete deacylation of d_5_-PG to GPG followed by CLN8-mediated reacylation with C22:6-CoA (Fig. 2A). As expected, CLN5 KO cells accumulated labeled C22:6-LPG with C22:6 esterified equally to the d_5_-labeled and unlabeled glycerol moieties (Fig. 2C). CLN5/6 DKO cells, by contrast, formed less d_5_-C22:6-LPG and lost this symmetry: species carrying C22:6 on the labeled glycerol were far more abundant than those carrying it on the unlabeled glycerol, which remained at background (Fig. 2C). Because CLN8 acylates the *S*-configured glycerol of GPG and is essentially unreactive towards *R,R*-GPG (Fig. 2D) (*8*, *13*), these data indicate that CLN6 converts the unlabeled *R*-glycerol to the *S* configuration; without CLN6 it remains *R*-configured and a poor CLN8 substrate, restricting acylation to the *S*-configured, d_5_-labeled glycerol.

To test this directly, we developed a two-step assay that reconstitutes consecutive reactions of the pathway and exploits the stereoselectivity of recombinant CLN8 for *S,S*-GPG (*8*). *R,S*- GPG was incubated with purified CLN6, after which CLN6 was heat-inactivated before adding CLN8 and C22:6-CoA. Thus, C22:6-LPG formation reports the product of the CLN6 reaction without requiring the two enzymes to act together (Fig. 2E, fig. S4A). Because CLN8 acylates *S,S*-GPG efficiently but *R,S*-GPG poorly, C22:6-LPG formation above the *R,S*-GPG baseline reports conversion to the *S,S* configuration during the first step. Initial C22:6-LPG production was limited until Ca^2+^ was included in the CLN6 incubation. In the presence of Ca^2+^, C22:6-LPG formation increased with incubation time and CLN6 concentration (Fig. 2F, fig. S4B). The apparent EC_50_ for Ca^2+^ was 1.3 mM, and a metal ion screen showed the reaction to be selective for Ca^2+^ among the cations tested (Fig. 2, G and H). In a separate binding assay, GPG, but not glycerophosphocholine or glycerophosphoethanolamine, showed affinity for Ca^2+^ in solution (fig. S4C). The reaction was also reversible: preincubating *S,S*-GPG with CLN6 reduced its subsequent conversion to C22:6-LPG by CLN8 (Fig. 2I). CLN6 also converted the non-physiological *R,R*-GPG into a CLN8-reactive product, although less efficiently than *R,S*-GPG, consistent with the two stereoinversions required to generate *S,S*-GPG (fig. S4D).

To establish the stereochemical outcome directly, we prepared a d_5_-*R,rac*-GPG tracer in which the deuterium label lies on the *R*-configured glycerol — opposite to the labeling in the cellular tracer. We then repeated the two-step assay, using LC-MS/MS fragmentation to determine which glycerol moiety received the C22:6 chain (Fig. 2J). Without CLN6, the reaction proceeded slowly and C22:6 was incorporated exclusively onto the unlabeled glycerol (Fig. 2K). Adding CLN6 in the first step strongly increased d_5_-C22:6-LPG formation and produced equal amounts of the two positional species, mirroring the symmetry seen in cells and demonstrating stereoconversion of the labeled *R*-glycerol by purified CLN6 (Fig. 2K). Finally, we asked whether CLN6 deficiency alters the stereochemistry of GPG within tissue lipids. Methanolysis of total liver lipids released GPG headgroups from PG, LPG and BMP, which were quantified and supplied in equal amounts to CLN8 (Fig. 2L, fig. S4E). Headgroups from *Cln6^nclf/nclf^* livers supported markedly less C22:6-LPG formation than those from wild-type livers, indicating depletion of *S,S*-configured lipids in CLN6-deficient liver (Fig. 2M). Together, these cellular and biochemical experiments identify CLN6 as a Ca^2+^-dependent GPG epimerase that converts *R,S*-GPG to *S,S*-GPG upstream of CLN8.

### Catalytic mechanism of CLN6

To investigate the biochemical mechanism of CLN6, we first used a high-confidence AlphaFold 3 model of the CLN6 homodimer to identify potential catalytic features (*27*). The model contained a central cavity (Fig. 3A). To determine how this cavity is oriented within the ER membrane, we combined localization and topology mapping. An HA-tagged CLN6 construct localized to the ER (fig. S5), and substituted cysteine accessibility experiments showed that the N terminus is cytosolic and the C terminus is luminal, placing the central cavity open to the ER lumen and agreeing with previous topology studies (fig. S6, A and B) (*28*). Blind molecular docking of *R,S*-GPG across the entire protein identified the highest-scoring pose within this lumen-facing cavity, positioning the predicted substrate-binding pocket in the Ca^2+^-rich ER lumen (Fig. 3A). To test the functional importance of this predicted substrate-binding pocket, we performed an alanine scan of residues surrounding the docked substrate (fig. S6C) and expressed each mutant in CLN6 KO HeLa cells. Although all mutant proteins were expressed at levels comparable to WT CLN6 and retained normal ER localization (fig. S6, D and E), D59A, D131A, D173A, E174A, H178A and E227A failed to restore 22:6-22:6 BMP in CLN6 KO cells, whereas the remaining mutants rescued this species to varying degrees (Fig. 3B). Together, these results identify a candidate catalytic center within the luminal cavity and residues required for CLN6 activity in cells.

**Fig. 3.**
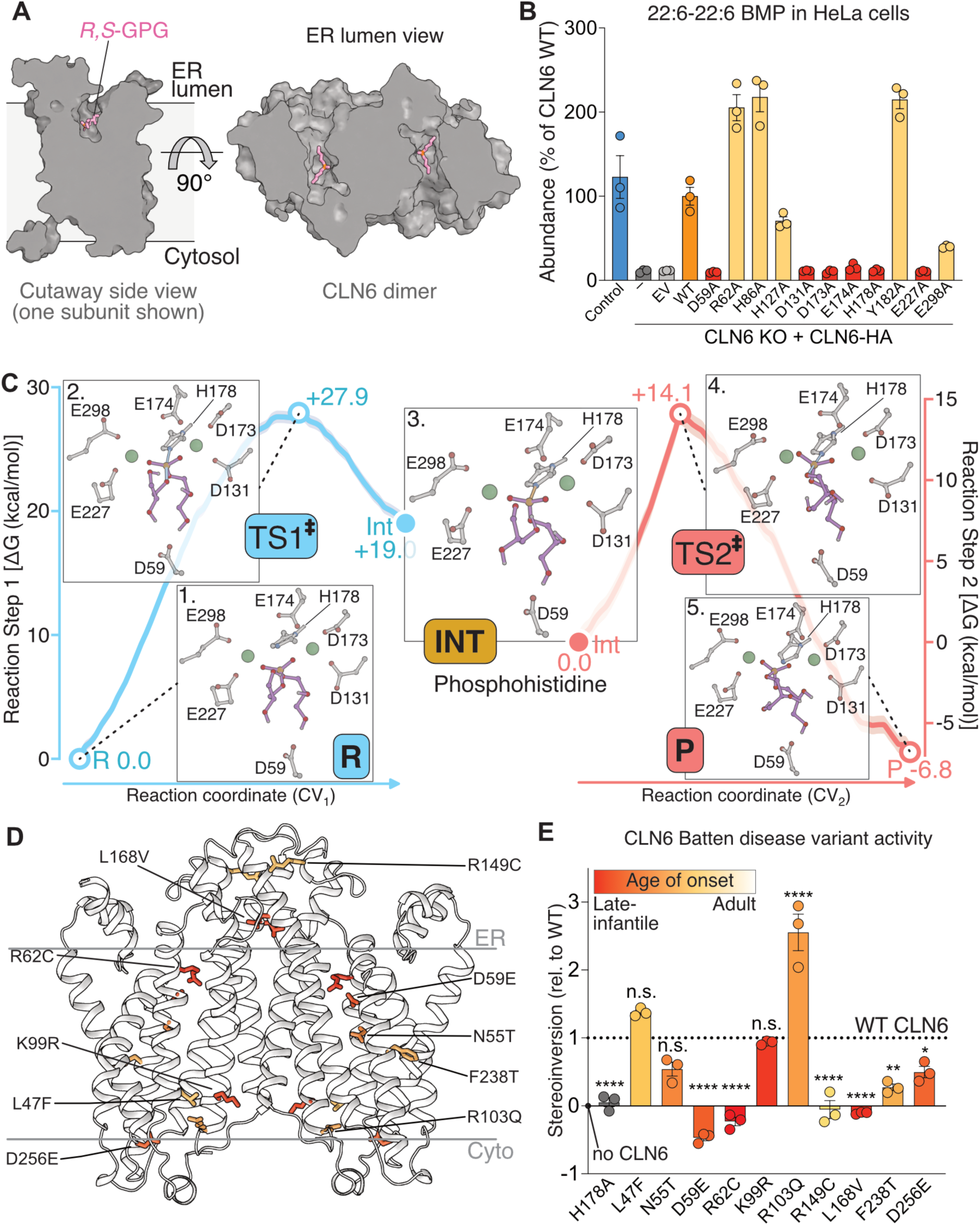
Catalytic mechanism of CLN6-mediated GPG stereoconversion and its perturbation in CLN6 Batten disease. (**A**) AlphaFold 3 model of the CLN6 homodimer with docked *R,S*-GPG (magenta). A cutaway side view of one subunit (left) and an ER lumen-facing view of the dimer (right) show the substrate-binding cavities. (**B**) Abundance of 22:6-22:6 BMP in control HeLa cells, CLN6 KO cells and CLN6 KO cells transfected with empty vector (EV) or HA-tagged WT CLN6 or the indicated alanine-substitution variants. BMP was measured by targeted LC-MS/MS and expressed relative to CLN6 KO cells expressing WT CLN6 (n = 3). (**C**) QM/MM free-energy profiles and representative structures for the proposed two-step His178-mediated phosphoryl-transfer reaction. *R,S*-GPG is initially oriented with its *S*-glycerol facing Asp59. His178 Nε2 attacks the substrate phosphorus to form a covalent phosphohistidine intermediate; a primary oxygen of the *R*-glycerol then attacks the phosphorus and displaces His178, yielding *S,S*-GPG. Structures show the reactant (R), transition states (TS1 and TS2), phosphohistidine intermediate (Int) and product (P); green spheres denote Ca^2+^ ions. Step 1 (blue, left axis) is referenced to R, whereas step 2 (red, right axis) is referenced to Int. (**D**) Positions of the analyzed Batten disease-associated substitutions in the CLN6 homodimer. Colors indicate the reported age at disease onset, from late-infantile (red) to adult-onset disease (yellow). (**E**) Activity of purified CLN6 disease-associated variants and the catalytically inactive H178A control, measured by C22:6-LPG formation in the two-step CLN6–CLN8 assay (n = 3 separate enzymatic reactions). Signals were corrected for the CLN8-only background and normalized to WT CLN6 activity (dotted line, 1). Negative values indicate product signals below this background. Variant colors correspond to those in (**D**). Data in (**B**) and (**E**) are means ± SEM. In (**E**), P < 0.05 (*), P < 0.01 (**) and P < 0.0001 (****); n.s., not significant, by one-way ANOVA with Dunnett’s multiple-comparisons test against WT CLN6. In vitro assay results are representative of two independent protein purifications.

Having identified candidate catalytic residues, we used structural and reaction modelling to investigate how CLN6 could convert *R,S*-GPG to *S,S*-GPG. We used Boltz-2 to co-fold CLN6 with *R,S*-GPG, specifying zero, one or two Ca^2+^ ions per active site and applying restraints to the region identified by mutagenesis (*29*). Comparison of protein–ligand interface confidence metrics guided the selection of a two-Ca^2+^ structure as a working model for molecular dynamics and quantum mechanics/molecular mechanics (QM/MM) calculations (fig. S7A). In this model, the substrate phosphate lies between the two Ca^2+^ ions in a carboxylate-rich cavity adjacent to His178 (fig. S7B).

We embedded the modelled CLN6 homodimer in an ER-like membrane and compared two substrate orientations using classical molecular dynamics. Each protomer contained *R,S*-GPG, with either the *R*-or *S*-glycerol moiety facing Asp59, a residue bordering the modelled glycerol-binding region (fig. S7B). During the unrestrained simulation, the orientation with the *S*-glycerol facing Asp59 allowed the free primary hydroxyl of the *R*-glycerol to fold back towards the phosphate, positioning it for a possible intramolecular phosphoryl transfer. To quantify this geometry, we counted frames in which both the hydroxyl oxygen and His178 Nε2 were within 4.5 Å of the phosphate phosphorus. This arrangement occurred in 8.6% of analyzed frames for this orientation and was not observed for the opposite orientation (fig. S7C). We selected a representative frame meeting these distance criteria as the starting structure for hybrid QM/MM calculations.

The QM/MM free-energy calculations supported a two-step phosphoryl-transfer mechanism mediated by a covalent phospho-His178 intermediate (Fig. 3C, movie S1). In the first step, His178 Nε2 attacks the substrate phosphorus as the original phosphoester bond breaks. The calculated free-energy surface supports a concerted, associative S_N_2-type substitution, without a resolved phosphorane or metaphosphate intermediate (fig. S8A). Concurrent proton transfer from the free primary hydroxyl of the *R*-glycerol to the departing oxygen generates the nucleophile for the second step. This deprotonated oxygen then attacks the phosphorus and displaces His178, producing *S,S*-GPG. Changes in atomic charge and the persistent, approximately six-coordinate geometry of both modelled Ca^2+^ ions were consistent with roles in charge stabilization and substrate organization during phosphoryl transfer (fig. S8, B and C). The proposed mechanism therefore converts the *R*-glycerol to the *S* configuration by transferring the phosphate linkage between its two primary hydroxyls, while preserving all bonds to the central chiral carbon (fig. S8D).

This reaction model is consistent with isotope-tracing evidence. In our in vitro CLN6 assay using d_5_-GPG as substrate, the d_5_ label was retained, with no d_4_-22:6-LPG detected, arguing against an oxidation–reduction mechanism involving loss of an *R*-glycerol-bound hydrogen (Fig. 2K). Furthermore, earlier work showed that labelling the terminal carbon distal to the phosphate in the *R*-glycerol moiety of *R,S*-PG resulted in this carbon becoming directly linked to the phosphate in BMP, consistent with migration of the phosphate to the opposite end of the glycerol (*19*). Together, these data support a direct intramolecular 1,3-phosphoryl migration in which the reacting *R*-glycerol moiety remains intact.

We next examined how disease-associated missense variants affect CLN6 enzymatic activity. We selected ten variants spanning different amino acid chemistries and positions within the CLN6 homodimer, and associated with clinical presentations ranging from severe late-infantile disease to the milder, adult-onset Kufs type A form (Fig. 3D) (*30–32*). All variants were expressed in cells, retained ER localization and were purified with yields and purity comparable to WT CLN6 (fig. S9, A and B). We compared their activities with WT CLN6 using our established in vitro assay (Fig. 2E). The catalytically inactive H178A control abolished CLN6-dependent stereoinversion, whereas the disease-associated variants displayed activities ranging from complete loss to approximately 2.5-fold WT activity (Fig. 3E). D59E and R62C produced less C22:6-LPG than the no-CLN6 control, yielding negative values after background subtraction, potentially reflecting reduced *R,S*-GPG availability for CLN8. Mapping these activities onto the CLN6 structural model revealed a spatial pattern: substitutions near the ER lumen-facing catalytic region, including D59E, R62C, R149C and L168V, abolished activity, whereas substitutions towards the cytosolic face generally retained partial, WT-like or enhanced activity (fig. S9C). Together, these findings support loss of enzymatic activity and consequent disruption of BMP synthesis as a molecular basis for the pathogenicity of several active-site-proximal CLN6 variants.

### CLN6-dependent BMP synthesis maintains lysosomal hydrolase abundance

Finally, we asked whether BMP deficiency underlies the lysosomal hydrolase depletion seen in CLN6 deficiency. This depletion has been attributed to a direct trafficking function, formalized in the ER-to-Golgi relaying of enzymes of the lysosomal system (EGRESS) model, in which a CLN6-CLN8 complex escorts soluble hydrolases from the ER to the Golgi (*5*, *33*). Our identification of CLN6 and CLN8 as biosynthetic enzymes suggested an alternative explanation: that loss of CLN6 disrupts BMP synthesis, which in turn causes lysosomal hydrolase depletion.

To test this hypothesis, we compared CLN6 KO HeLa cells with isogenic control cells by whole-cell proteomics. CLN6 KO cells showed reduced abundance of multiple lysosomal hydrolases, including acid sphingomyelinase (ASM; encoded by *SMPD1*), CTSD, PPT1 and TPP1 (Fig. 4A). We therefore used these four proteins as markers to determine whether restoring BMP synthesis could correct the lysosomal hydrolase defect.

**Fig. 4.**
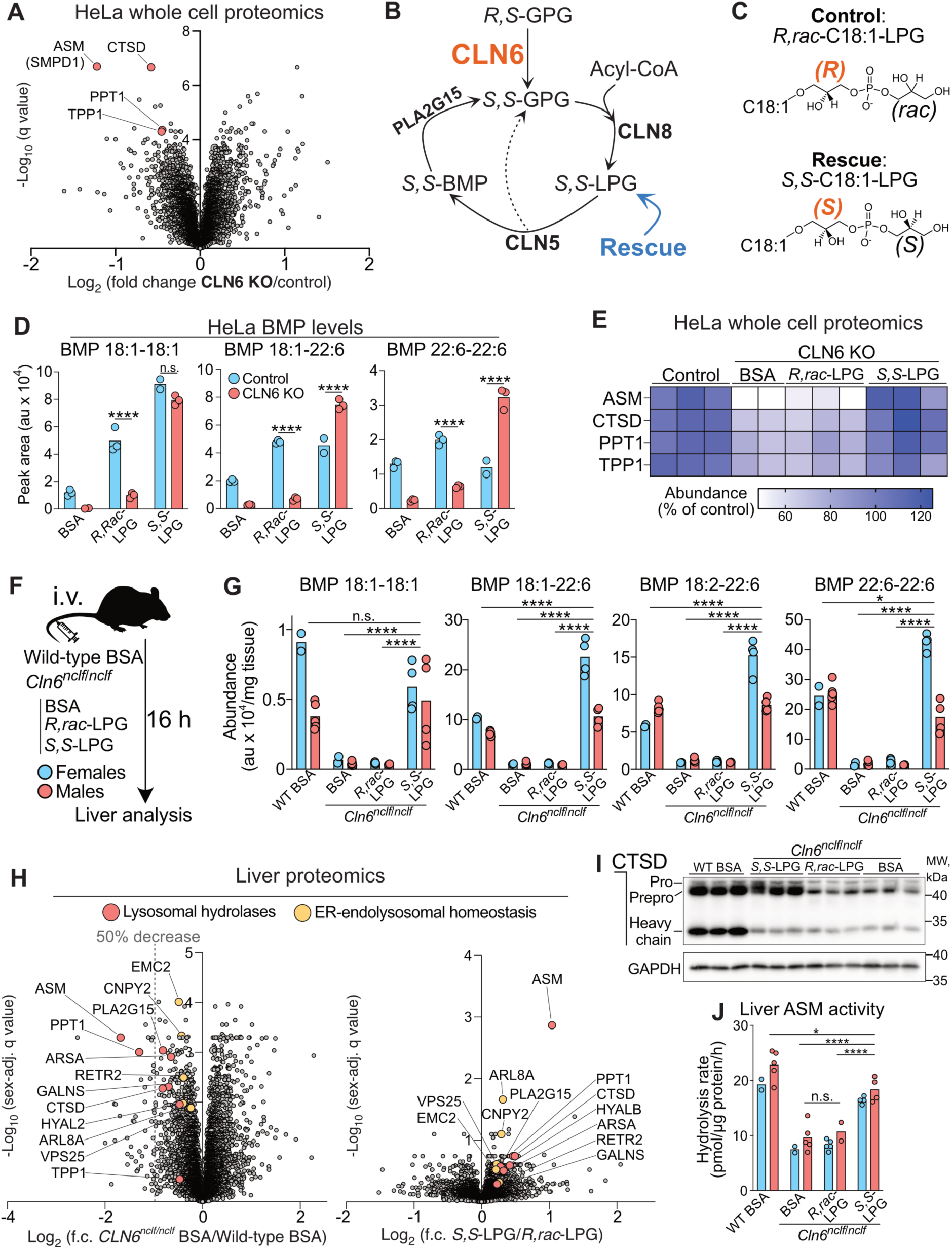
Restoring BMP reverses lysosomal hydrolase depletion in CLN6-deficient models. (**A**) Whole-cell proteomic comparison of CLN6 KO and control HeLa cells (n = 6 biological replicates per group), with selected lysosomal hydrolases highlighted in red. (**B**) BMP biosynthetic and recycling pathway, showing CLN6-dependent formation of *S,S*-GPG and the point at which *S,S*-LPG supplementation bypasses CLN6. (**C**) Structures of *R,rac*-C18:1-LPG (control) and *S,S*-C18:1-LPG (rescue). (**D**) Abundance of the indicated BMP species in control and CLN6 KO HeLa cells treated for 16 h with BSA or 10 μM *R,rac*-LPG or *S,S*-LPG (n = 3). (**E**) Heatmap of lysosomal hydrolase abundance measured by whole-cell proteomics after 48 h of the indicated treatments. Each column represents one biological replicate, with abundance expressed as a percentage of the mean in BSA-treated control cells (n = 2 for BSA-treated CLN6 KO cells; n = 3 for all other groups). (**F**) Mouse treatment protocol. Wild-type mice received BSA, and *Cln6^nclf/nclf^*mice received BSA or a single intravenous dose of 2 μmol BSA-complexed *R,rac*-C18:1-LPG or *S,S*-C18:1-LPG. Livers were collected 16 h later. (**G**) Hepatic abundance of the indicated BMP species in 7- to 8-week-old littermates. Groups comprised BSA-treated wild-type mice (n = 7; 2 female, 5 male), BSA-treated *Cln6^nclf/nclf^*(n = 7; 2 female, 5 male), and *Cln6^nclf/nclf^* mice treated with *R,rac*-LPG (n = 7; 5 female, 2 male) or *S,S*-LPG (n = 8; 4 female, 4 male). The same groups were analyzed in (**H**) and (**J**). BMP in (**D**) and (**G**) was measured by targeted LC-MS/MS. (**H**) Liver proteomic comparisons of BSA-treated *Cln6^nclf/nclf^* versus wild-type mice (left) and *S,S*-LPG-versus *R,rac*-LPG-treated *Cln6^nclf/nclf^*mice (right). Fold changes and statistical tests were adjusted for sex. Selected lysosomal hydrolases are highlighted in red and proteins associated with ER–endolysosomal homeostasis in yellow. (**I**) Representative immunoblot of CTSD in liver lysates from the indicated groups, with GAPDH as a loading control. Immunoblots from the remaining mice are shown in fig. S10F. (**J**) Hepatic acid sphingomyelinase (ASM; encoded by *SMPD1*) activity, expressed as substrate hydrolysis per μg protein per hour. In (**D**), (**G**) and (**J**), points represent individual samples and bars indicate means; females and males are shown separately in blue and red in (**G**) and (**J**). In (**A**) and (**H**), q values were calculated using Benjamini–Hochberg correction across all tested proteins. Statistical comparisons in (**D**) used two-way ANOVA with Šídák’s multiple-comparisons test; those in (**G**) and (**J**) used two-way ANOVA with experimental group and sex as factors, followed by Tukey’s multiple-comparisons test. *P* < 0.05 (*), *P* < 0.001 (***) and *P* < 0.0001 (****); n.s., not significant.

We reasoned that CLN6 deficiency should be particularly amenable to metabolic rescue by an *S,S*-configured intermediate because CLN6 establishes the *S,S* configuration at the point of entry into the BMP biosynthetic cycle (Fig. 4B). Once an *S,S*-configured intermediate has been supplied, the *S,S*-GPG backbone can therefore undergo repeated rounds of acylation, BMP formation and degradation without requiring further CLN6 activity (Fig. 4B).

We first tested whether exogenous GPG could restore BMP synthesis in CLN6 KO cells. A *rac,rac*-[^13^C_5_]GPG stereoisomer mixture generated ^13^C_5_-labeled LPG and BMP, showing that exogenous GPG is taken up and that its *S,S* isomer enters the pathway without CLN6 (fig. S10, A to C). The rescue was stereospecific but partial: total BMP rose only modestly after *rac,rac*-GPG treatment, remaining well below control levels, whereas *R,rac*-GPG, which lacks the *S,S* isomer, had no effect; this limited restoration is consistent with the poor cellular uptake of GPG observed previously (*13*).

We therefore turned to *S,S*-C18:1-LPG, which CLN5 can use to produce 18:1-18:1 BMP while releasing *S,S*-GPG (Fig. 4C). In our previous study, *S,S*-C18:1-LPG restored 18:1-18:1 BMP in CLN8 KO cells and mice, but the rescue was restricted to this species because the released *S,S*-GPG could not be reacylated in the absence of CLN8 (*8*). CLN6 KO cells, by contrast, retain active CLN8, allowing CLN5-generated *S,S*-GPG to be reacylated with endogenous fatty acyl-CoAs and re-enter the BMP biosynthetic cycle (Fig. 4B). We therefore predicted that *S,S*-C18:1-LPG would restore a broader range of BMP species in CLN6 KO cells than previously observed in CLN8 KO cells. Consistent with this model, treatment with *S,S*-C18:1-LPG for 16 h restored all measured BMP species to control levels, whereas *R,rac*-C18:1-LPG did not (Fig. 4D). Both LPG stereoisomers increased cellular C18:1-LPG abundance well above BSA-treated controls, indicating comparable uptake (fig. S10D). To determine whether BMP restoration also corrected lysosomal hydrolase abundance, we treated CLN6 KO cells separately with either LPG stereoisomer for 48 h and performed whole-cell proteomics. *S,S*-LPG restored ASM, CTSD, PPT1 and TPP1 abundance to control levels, whereas *R,rac*-LPG had no effect (Fig. 4E). The stereospecific restoration of both BMP and lysosomal hydrolases demonstrated that defective BMP synthesis causes lysosomal hydrolase depletion in CLN6 KO cells.

Last, we asked whether BMP deficiency and lysosomal hydrolase depletion could also be reversed in vivo. We previously established that intravenously administered *S,S*-C18:1-LPG rapidly reaches the liver but does not cross the blood-brain barrier in mice; we therefore assessed rescue in the liver (*8*). *Cln6^nclf/nclf^*mice were injected intravenously with *R,rac*- or *S,S*-C18:1-LPG, and their livers were collected 16 h later, alongside BSA-treated wild-type and *Cln6^nclf/nclf^* littermates (Fig. 4F). *S,S*-LPG restored all measured hepatic BMP species in *Cln6^nclf/nclf^*mice to the levels observed in BSA-treated WT mice, whereas hepatic BMP remained depleted after *R,rac*-LPG treatment (Fig. 4G). The altered fragmentation of asymmetric BMP species seen in untreated mutant tissue was also corrected by *S,S*-LPG, but not by *R,rac*-LPG (fig. S10E). Whole-liver proteomics showed that BSA-treated *Cln6^nclf/nclf^* mice were deficient in numerous lysosomal enzymes, with ASM, CTSD and PPT1 among the most strongly depleted, and a single acute dose of *S,S*-LPG increased all three relative to *R,rac*-LPG-treated *Cln6^nclf/nclf^* mice (Fig. 4H). The increase in ASM remained significant after proteome-wide multiple-testing correction (*q* = 0.0013, *P* = 1.8 × 10^-7^), whereas the increases in CTSD and PPT1 were significant before, but not after, correction (CTSD, *q* = 0.18, *P* = 0.00053; PPT1, *q* = 0.18, *P* = 0.00044) (Fig. 4H). Immunoblotting supported increased CTSD abundance, with a more pronounced recovery of the precursor form than of the mature heavy chain after 16 h (Fig. 4I, fig. S10F). The increase in hydrolase abundance was accompanied by increased hepatic ASM activity after *S,S*-LPG treatment (Fig. 4J).

Together, these experiments establish defective BMP synthesis as the cause of lysosomal hydrolase depletion in CLN6 Batten disease. A single dose of *S,S*-LPG normalized BMP levels and increased lysosomal hydrolase abundance and activity in vivo, demonstrating that this hydrolase depletion is a rapidly reversible consequence of BMP deficiency.

## Discussion

Here we identify CLN6 as a Ca^2+^-dependent GPG epimerase that establishes the unusual *S,S* stereochemistry of BMP. Together with CLN8 and CLN5, CLN6 defines an enzymatic sequence from *R,S*-GPG to *S,S*-BMP, resolving the previously unassigned stereochemical step (fig. S11). These findings place three sequence-unrelated Batten disease proteins in a shared biosynthetic pathway and provide a biochemical basis for their overlapping disease phenotypes. An independent study identifies the same CLN6 activity using complementary stereochemical analyses and demonstrates accumulation of *S,R*-LPG and *R,S*-BMP in its absence (*34*). These stereochemical changes are consistent with the altered fragmentation of residual BMP observed here in CLN6-deficient tissues and with our observation that CLN8 acylates *R,S*-GPG exclusively at the *S*-configured glycerol moiety in vitro. CLN6’s position upstream of CLN8 and its Ca^2+^ dependence suggest that it could also regulate the flux of GPG into the BMP pathway, coupling BMP synthesis to cellular stimuli that alter ER luminal Ca^2+^ availability.

Our findings also establish a physiological role for BMP in maintaining lysosomal hydrolase abundance. CLN6 deficiency depleted multiple lysosomal enzymes, including PPT1, TPP1 and CTSD, whose individual deficiencies cause other forms of Batten disease (*1*). Restoration of hydrolase abundance while CLN6 remains absent shows that this defect can be reversed by bypassing the CLN6-dependent step in BMP synthesis. Together with the BMP deficiency reported in CLN3, CLN5, CLN8 and CLN11 disease (*8*, *10*, *11*, *22*, *35*), these findings support a unifying mechanism across Batten disease: loss of lysosomal hydrolase function can arise either directly from mutations in the enzymes themselves or indirectly through BMP deficiency that depletes multiple hydrolases. Whether BMP maintains hydrolase abundance by controlling enzyme delivery, maturation or stability remains to be determined.

These findings support BMP restoration as a therapeutic strategy for CLN6 Batten disease and other BMP-deficient forms, with interventions tailored to the biosynthetic step disrupted by the affected *CLN* gene (fig. S11). Achieving neurological benefit will require effective central nervous system delivery, as *S,S*-C18:1-LPG does not cross the blood–brain barrier. More broadly, increasing lysosomal BMP abundance could improve lysosomal degradative capacity in common neurodegenerative disorders characterized by impaired lysosomal catabolism (*36*).

## Acknowledgments

We thank David Ron, Julian Rayner and Jonathon Nixon-Abell for advice; the Cambridge Institute for Medical Research Proteomics Core Facility and the CRUK Cambridge Institute Proteomics Core Facility for proteomic analyses; and the MRC Mitochondrial Biology Unit Light Microscopy Facility for microscopy support. We thank the patients and their families, and the healthy volunteers, for their contributions to this study. *R,S*-GPG and *S,S*-C18:1-LPG were synthesised by o2h Group and WuXi AppTec, respectively. Gene synthesis was performed by GenScript, Sanger sequencing by Source Genomics and mouse genotyping by Transgenix. ChatGPT (OpenAI) was used to assist with manuscript preparation.

## Funding

UKRI Medical Research Council grant MC_UU_00028 (PKS, KT, KP)

UKRI Medical Research Council grant MC_UU_00028/2 (ERSK)

UKRI Medical Research Council grant MC_UU_00028/4 (AMJ, CSY, MPM)

UKRI Medical Research Council grant MC_UU_00028/5 (JG, JP)

Academy of Medical Sciences Springboard Award SBF0010/1078 (KP)

Batten Disease Global Research Initiative award (KP)

NCL Foundation (NCL-Stiftung) award (KP)

UKRI Biotechnology and Biological Sciences Research Council grant BB/Y002865/1 (DL)

Wellcome Trust Investigator Award 220257/Z/20/Z (MPM)

German Federal Ministry of Research, Technology and Space (BMFTR) grant 01GL2404A (CP, AS)

Wellcome Trust grant 226800/Z/22/Z (BJ, AK)

NIHR Cambridge Biomedical Research Centre grant NIHR203312 (BJ, AK)

Austrian Science Fund (FWF) grant 10.55776/J4932 (DB)

Austrian Science Fund (FWF) grant 10.55776/F73 (RZ, KG)

French National Research Agency, France 2030 program, grant ANR-10-IAHU-01 (JP)

Novo Nordisk Foundation grant NNF24OC009614 (MJ)

Neurogene Inc. (DNP, NLM)

## Author contributions

Conceptualization: KP. Methodology: PKS, AMJ, ASS, JB, DL, DB, BJ, HEJ, SOL, CSY, KP. Investigation: PKS, AMJ, CP, ASS, JB, JG, MKP, SKL, KT, DL, DB, JVDK, MKB, KP. Formal analysis: AMJ, ASS, JB, JG, DL, BJ, HEJ, SOL, CSY, KP. Resources: CP, CDB, JMW, DNP, NLM, AS. Funding acquisition: JMW, AK, MJ, ERSK, DNP, NLM, JP, RZ, KG, AS, MPM, KP. Supervision: JMW, AK, MJ, ERSK, DNP, NLM, JP, RZ, KG, AS, MPM, KP. Writing – original draft: KP. Writing – review & editing: all authors.

## Data, code and materials availability

All data supporting the findings of this study are included in the manuscript and supplementary materials. The proteomics and untargeted lipidomics datasets will be deposited in public repositories, with accession codes provided in the published article. Processed lipidomics and proteomics datasets will also be included as supplementary tables. Requests for reagents or biological materials should be directed to Kasparas Petkevicius.

## Supplementary materials

## Materials and Methods

Figs. S1 to S11

Movie S1

Methods references (1–27)

## Materials and methods

### Reagents, antibodies and plasmids

Antibodies, reagents and plasmids are listed in the tables below and, where relevant, are also identified in the corresponding methods sections.

Antibodies used in this study

| Target | Supplier | RRID | Catalogue No. |
| --- | --- | --- | --- |
| Anti-CLN5 | Abcam | RRID:AB_3662651 | Ab170899 |
| Anti-CLN6 | Proteintech | RRID:AB_2878671 | 20315-1-AP |
| Anti-TMEM192 (EPR14330-67; BSA- and azide-free) | Abcam | RRID:AB_3095638 | ab232600 |
| Anti-Vinculin | Merck | RRID:AB_477629 | V9131 |
| Anti-FLAG M2 | Sigma | RRID:AB_262044 | F1804 |
| Anti-Lamin B1 | Invitrogen | RRID:AB_2281281 | MA1-06103 |
| Anti-HA tag | ThermoFisher | RRID:AB_2533988 | 71-5500 |
| Anti-SERCA2 ATPase | Abcam | RRID:AB_303961 | ab3625 |
| Anti-CTSD | Abcam | RRID:AB_1310298 | AB75852 |
| Anti-GAPDH | Proteintech | RRID:AB_2107436 | 60004-1-Ig |
| Anti-mouse IgG-HRP | Cell Signaling Technology | RRID:AB_330924 | 7076-S |
| Anti-rabbit IgG-HRP | Cell Signaling Technology | RRID:AB_2099233 | 7074-S |
| Anti-HA tag (C29F4; immunofluorescence) | Cell Signaling Technology | RRID:AB_1549585 | 3724 |
| Anti-Calnexin (W17077C) | BioLegend | RRID:AB_2728519 | 699401 |
| Anti-GM130 | BD Transduction Laboratories | RRID:AB_398141 | 610822 |
| F(ab')<sub>2</sub>-goat anti-rabbit IgG (H+L) cross-adsorbed, Alexa Fluor 488 conjugate | Invitrogen | RRID:AB_2534114 | A-11070 |
| Goat anti-mouse IgG1 cross-adsorbed, Alexa Fluor 568 conjugate | Invitrogen | RRID:AB_2535766 | A-21124 |
| Goat anti-rat IgG (H+L), Alexa Fluor 647 conjugate | Invitrogen | RRID:AB_141778 | A-21247 |

Chemicals and reagents used in this study

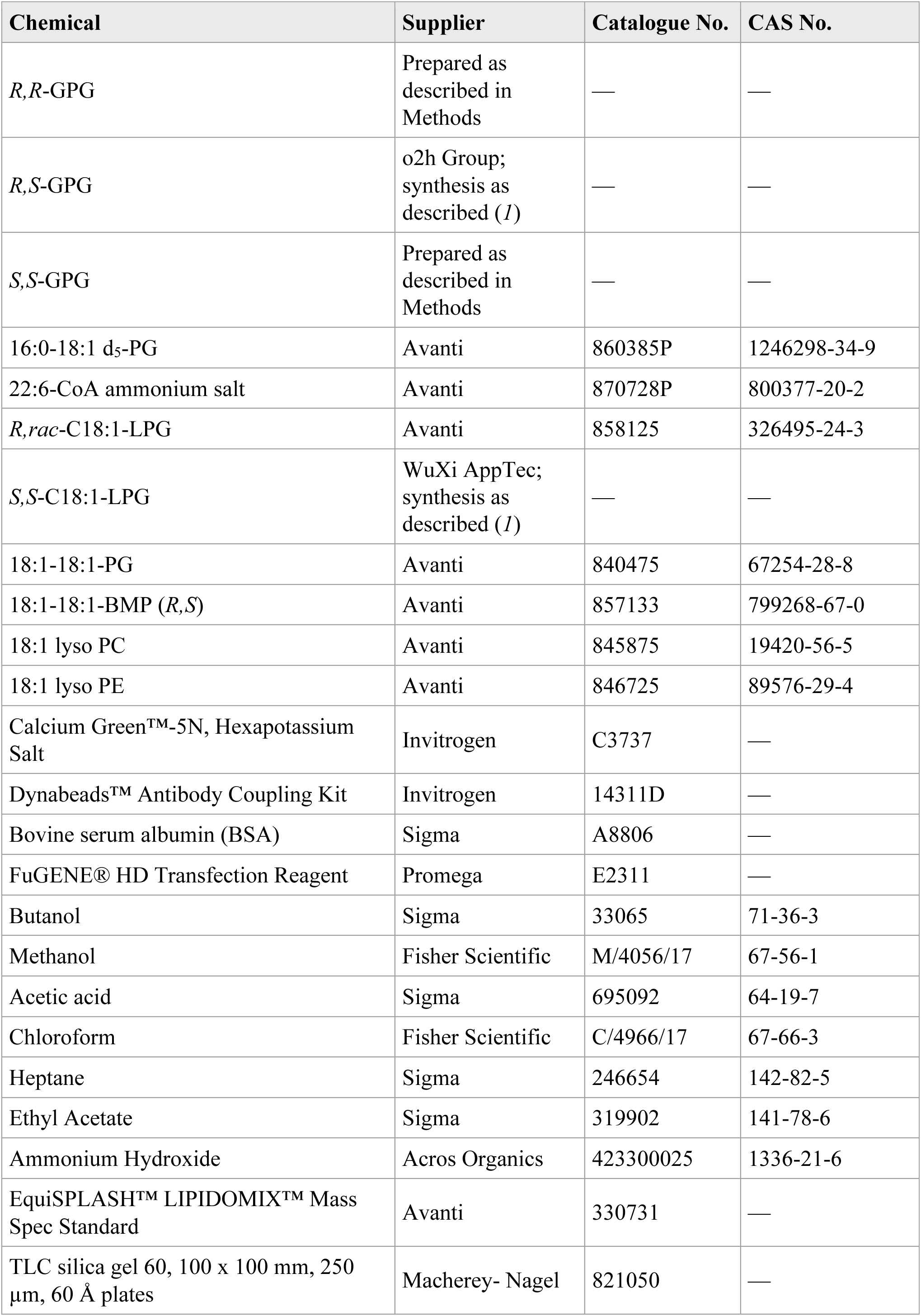
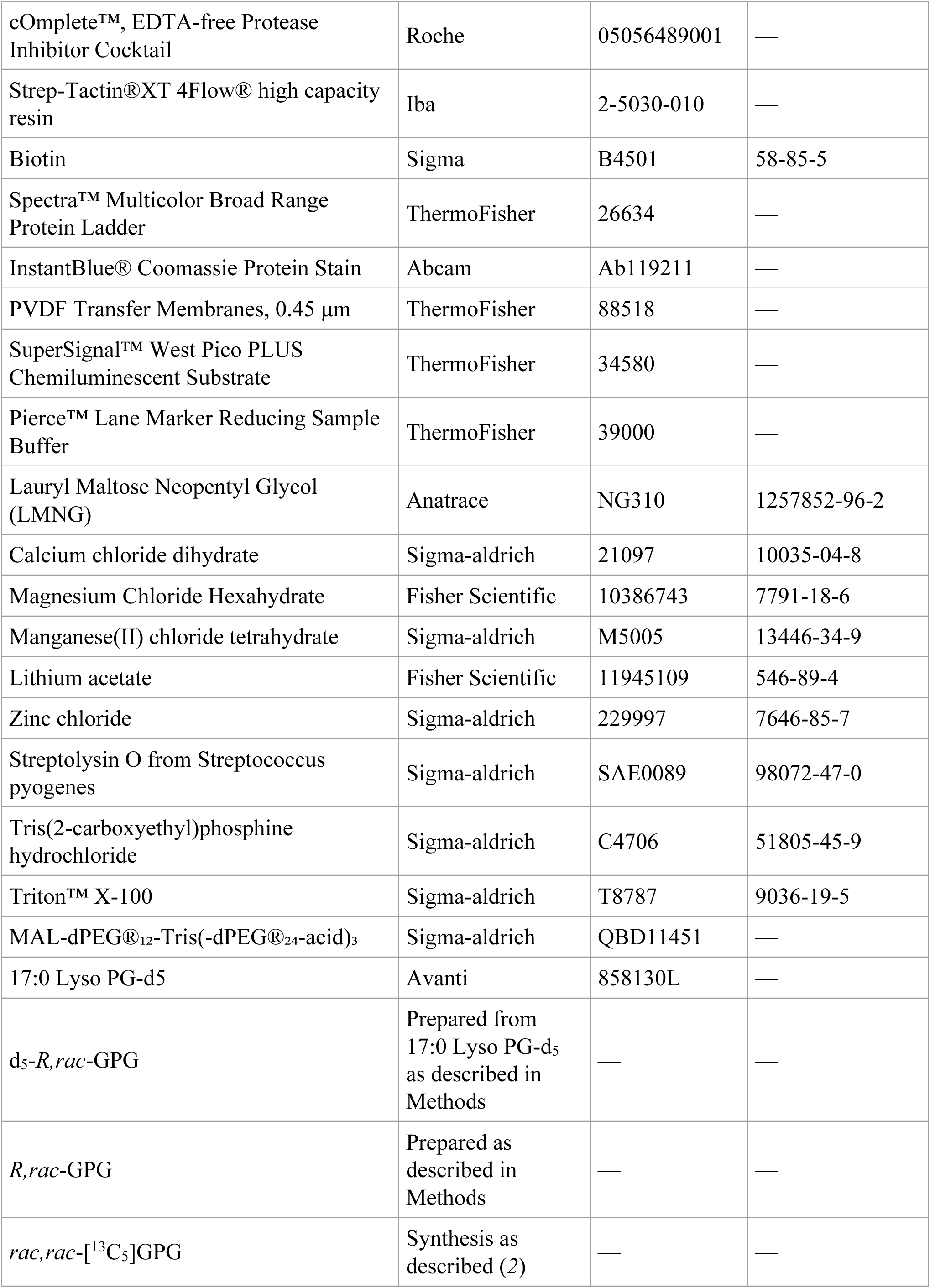
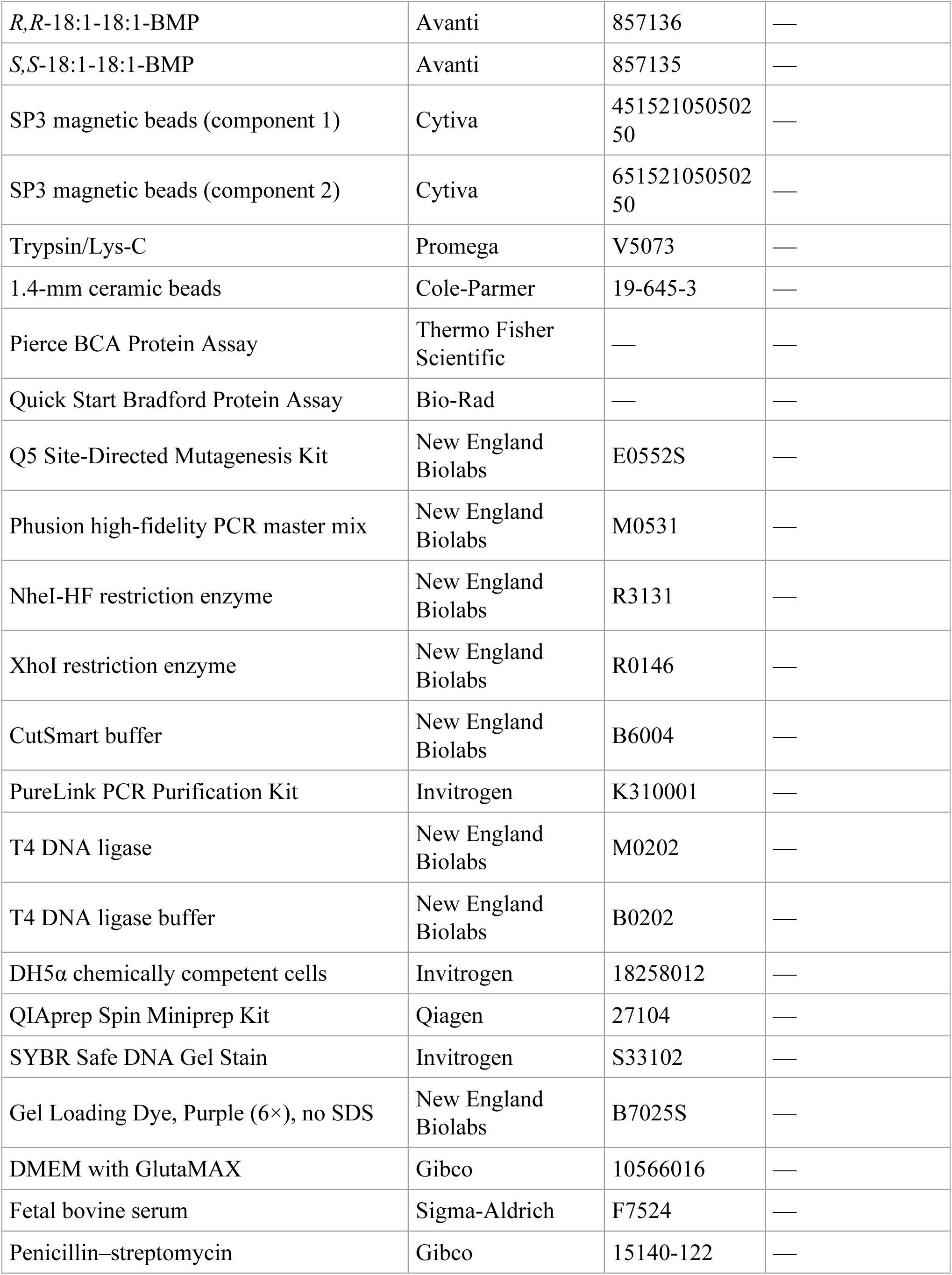
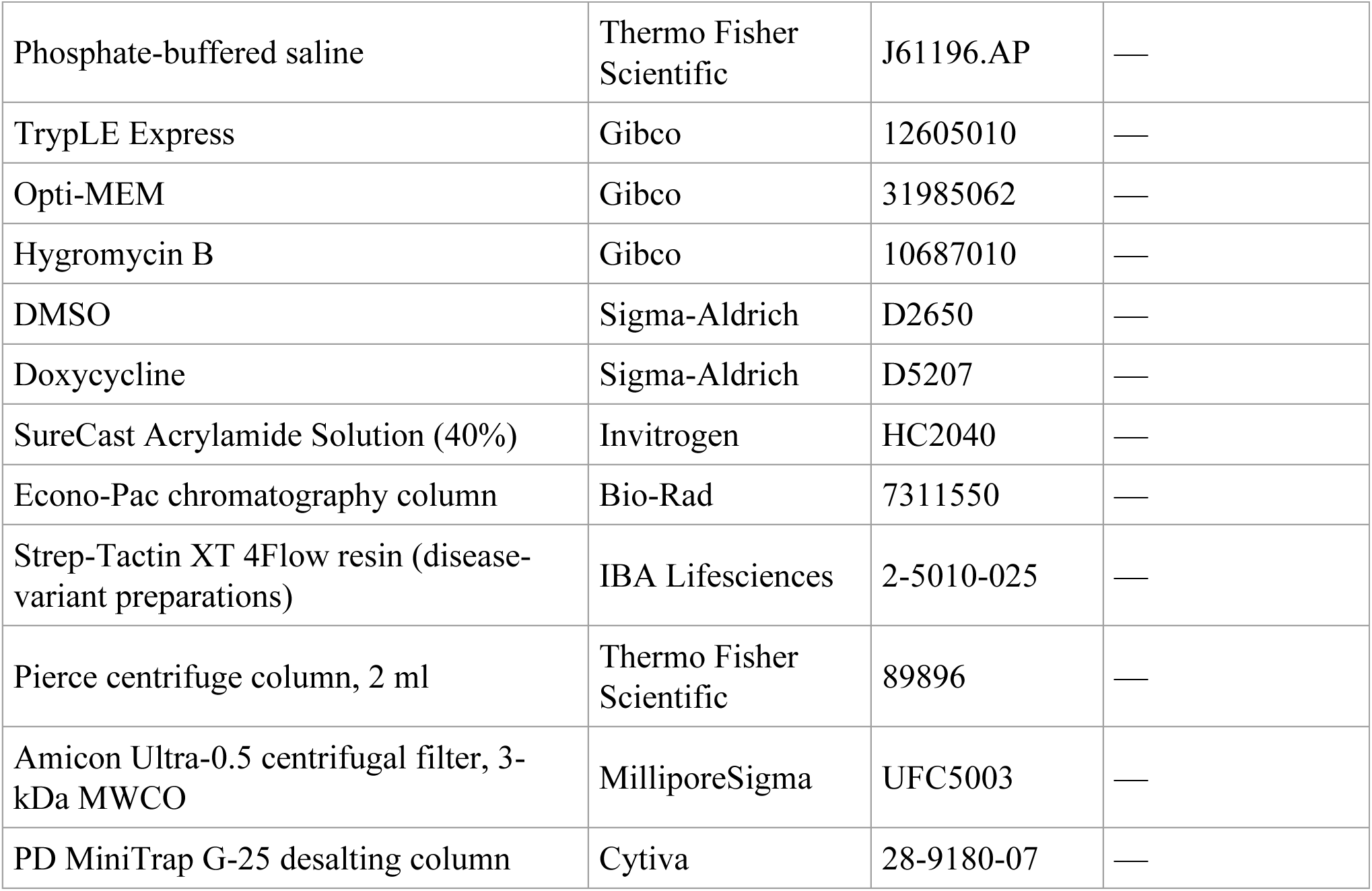

Plasmids used in this study

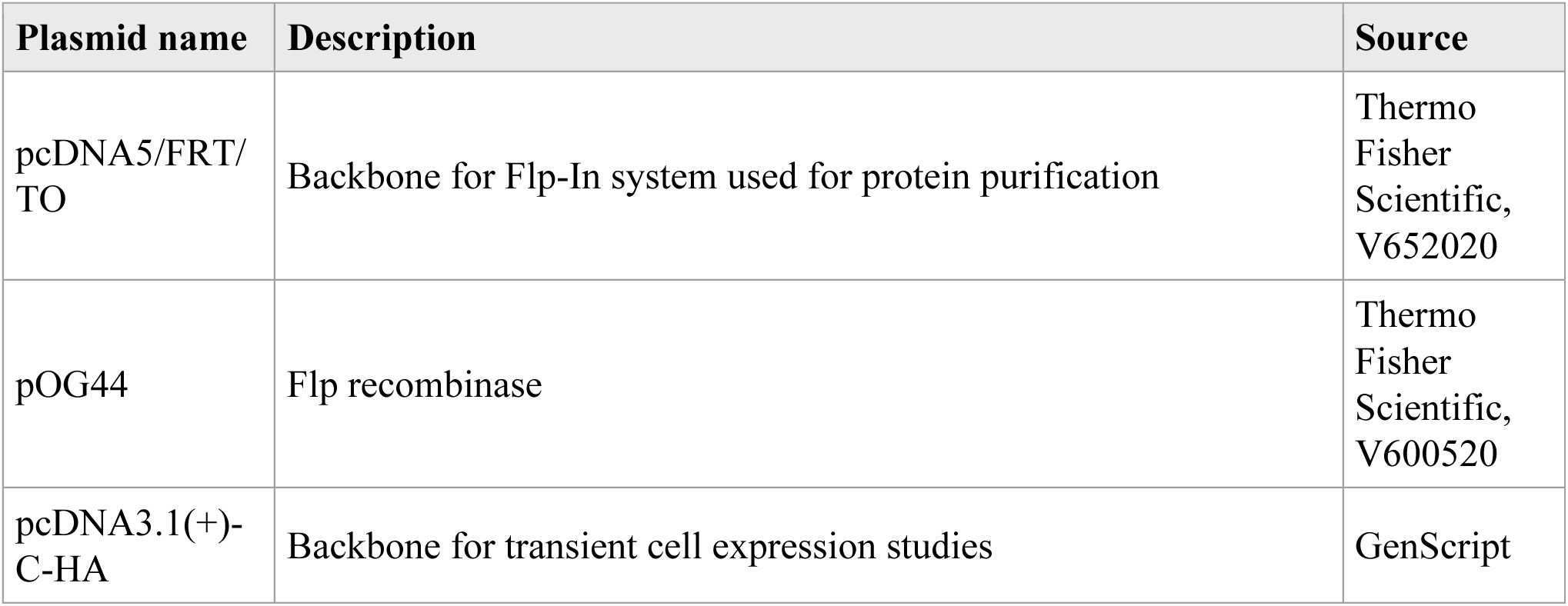
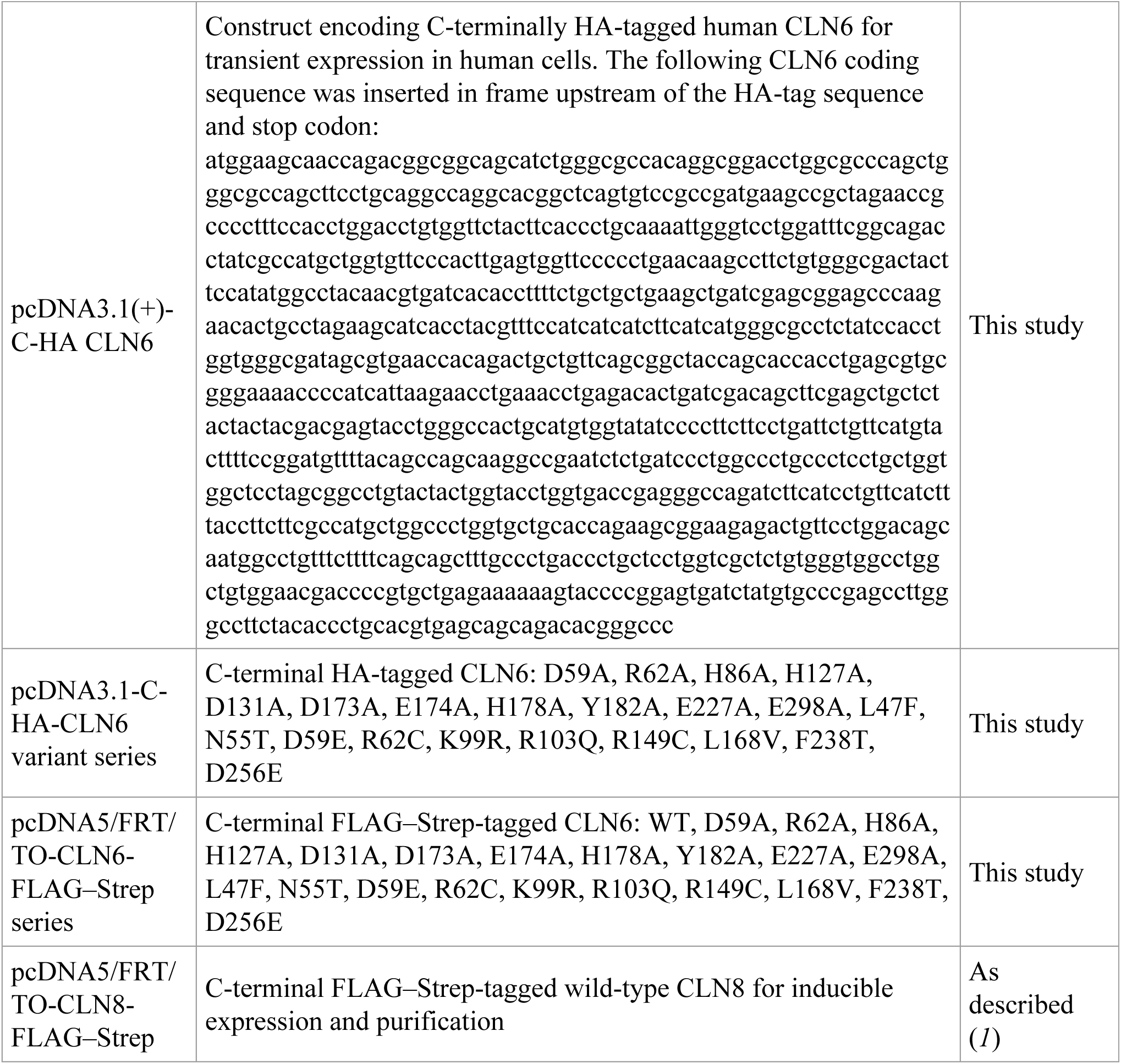

### Human participants

Three children with genetically confirmed CLN6 disease were recruited from the NCL specialty clinic at the University Children’s Hospital, University Medical Center Hamburg-Eppendorf, Hamburg, Germany. Three healthy adult donors were recruited at the same clinical site. Fresh peripheral blood (15 ml per participant) was collected into EDTA tubes. Participant age, sex, phenotype, age at symptom onset and CLN6 genotype are provided in the table below. The study was approved by the local medical ethics committee of the Ärztekammer Hamburg (PV7199). Written informed consent was obtained from patients or their parents in accordance with the Declaration of Helsinki.

#### Characteristics of participants providing peripheral blood samples

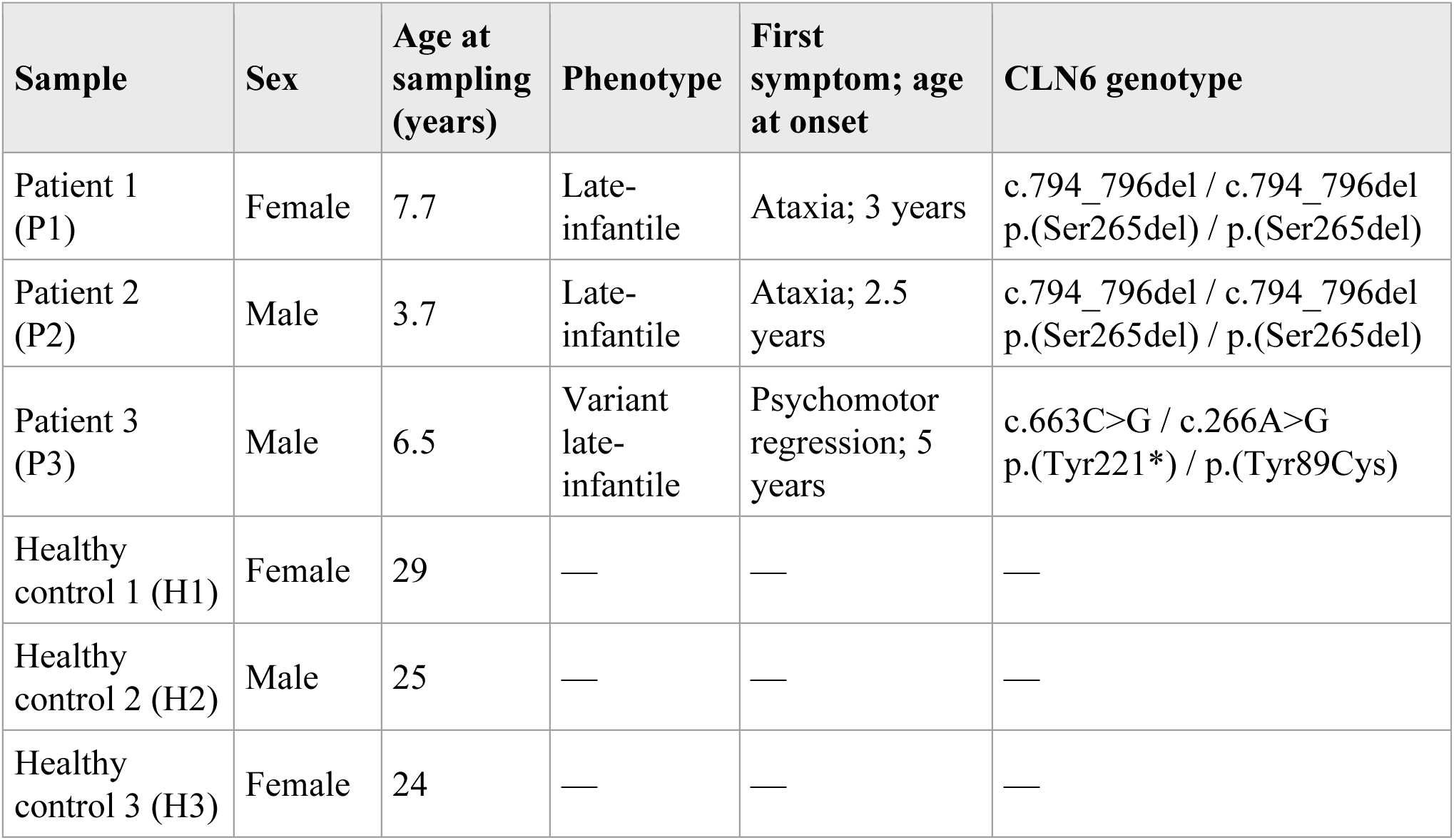

### Tagless lysosome immunoprecipitation from peripheral blood mononuclear cells

Tagless lysosome immunoprecipitation (LysoIP) was adapted from a recently published protocol (*3*). First, magnetic beads were coupled to an antibody recognizing the cytosolic domain of endogenous TMEM192 (rabbit monoclonal EPR14330-67, supplied BSA- and azide-free for coupling; Abcam, ab232600). 5 mg Dynabeads M-270 Epoxy were washed once with 1 ml C1 buffer from the Dynabeads Antibody Coupling Kit and collected on a magnetic rack for 1 min. The beads were resuspended in 100 µl anti-TMEM192 antibody (approximately 1 µg/µl) and 150 µl C1 buffer, followed by the addition of 250 µl C2 buffer. Antibody coupling was performed for 24 h at 37 °C with agitation at 300 rpm. The beads were subsequently washed once with HB buffer, once with LB buffer and twice with SB buffer, including a 15-min incubation during the final SB wash. Coupled beads were resuspended in 500 µl SB buffer and stored at 4 °C. This procedure provided an antibody loading of approximately 20 µg per milligram of beads.

Peripheral blood mononuclear cells (PBMCs) were isolated from 15 ml fresh EDTA-anticoagulated blood. Blood was diluted 1:1 with PBS at room temperature, layered into SepMate-50 tubes containing approximately 13 ml Ficoll-Paque PREMIUM and centrifuged at 1200 × g for 10 min at room temperature. The PBMC-containing layer was transferred to a fresh 50-ml tube, adjusted to 50 ml with PBS and centrifuged at 450 × g for 5 min. The resulting pellet was washed once more with 50 ml PBS under the same centrifugation conditions and resuspended in 1 ml ice-cold KPBS (136 mM KCl and 10 mM KH2PO4, adjusted to pH 7.25 with KOH). Immediately before use, KPBS was supplemented with one cOmplete protease-inhibitor tablet and one PhosSTOP phosphatase-inhibitor tablet per 10 ml. For each isolation, 100 µl TMEM192-coupled bead suspension was transferred to a protein-low-binding tube, collected magnetically for 1 min and washed with 1 ml inhibitor-supplemented KPBS. The PBMC suspension was centrifuged at 1,000 × g for 2 min at 4 °C, and comparable cell pellets were resuspended in 1 ml inhibitor-supplemented KPBS. Diisopropyl fluorophosphate (DIFP) was added to a final concentration of 0.5 mM, and 50 µl of the cell suspension was retained as the whole-cell input. A 0.5 M DIFP stock was prepared in isopropanol and stored at -80 °C. DIFP was handled in a fume hood, and contaminated waste was inactivated using 2% NaOH.

An Isobiotec Cell-Breaker fitted with a ball bearing providing a 12-µm clearance was pre-cooled on ice and equilibrated with 3 ml ice-cold inhibitor-supplemented KPBS. PBMCs were disrupted by passing the suspension through the chamber 20 times using two 1-ml syringes. The homogenate was collected in a protein-low-binding tube and centrifuged at 1,000 × g for 2 min at 4 °C. The resulting post-nuclear supernatant was incubated with 100 µl pre-washed TMEM192-coupled beads for 5 min at 4 °C with continuous rotation. Beads were captured on a magnetic rack for 60 s, and the unbound material was retained as the flow-through fraction. The bead-bound lysosome fraction was washed twice with 1 ml KPBS lacking inhibitors, resuspended in a further 1 ml KPBS and transferred to a fresh protein-low-binding tube before a final magnetic capture. Lysosome-bound bead fractions were stored at - 80 °C until lipid extraction.

### Ovine studies

Control, CLN5-deficient and CLN6-deficient sheep (*4*) brain tissues were supplied by Lincoln University, New Zealand. The cohort comprised five unaffected control female sheep, two homozygous CLN5-affected Borderdale female sheep, and two homozygous CLN6-affected South Hampshire female sheep. Animals were housed at Lincoln University under standard research husbandry conditions with ad libitum access to feed and water. Sheep were maintained in accordance with the New Zealand Animal Welfare Act (1999) and institutional animal welfare guidelines. All procedures were approved by the Lincoln University Animal Ethics Committee (AEC#2020-22 and AEC#2023-67), and the study was reported in accordance with ARRIVE 2.0 guidelines.

Animals were euthanized at 12 months of age, and brain tissue was collected immediately post-mortem. Samples from the cerebellum, thalamus, motor cortex and visual cortex were dissected, snap-frozen, stored at −80 °C, and transported on dry ice to the University of Cambridge for analysis.

### Mouse studies

Tissues analyzed in Figs. 1 and 2 and the corresponding supplementary figures were obtained from 6-month-old female wild-type and homozygous *Cln6^nclf/nclf^* mice at Sanford Research (Sioux Falls, SD, USA). Animals were bred, maintained and euthanized in an AAALAC International-accredited facility (USDA license 46-R-0009) under Sanford Research Institutional Animal Care and Use Committee approval (protocol 2023-0102), following institutional, federal and international animal-welfare guidelines. Mice were housed in groups of up to five in individually ventilated Tecniplast Green Line cages with HEPA-filtered airflow, under a 14-h light–10-h dark cycle at 24 ± 1 °C and 30–70% relative humidity. Irradiated Teklad 2018 chow and non-acidified water were available ad libitum; Teklad 7092–7097 corncob bedding and Teklad 7099 bedding enrichment were provided.

The LPG rescue experiment reported in Fig. 4, including intravenous administration and liver collection, was performed at the University of Cambridge under the UK Animals (Scientific Procedures) Act 1986 (Home Office project license PP1740969), following approval by the University’s Animal Welfare and Ethical Review Body and in accordance with EU Directive 2010/63/EU. Experiments are reported in accordance with ARRIVE 2.0 guidelines. The *Cln6^nclf^* line was obtained from The Jackson Laboratory (strain no. 002648; RRID: IMSR_JAX:002648). Homozygous *Cln6^nclf/nclf^* mice and wild-type littermates were generated by intercrossing heterozygous parents, and both sexes were used. Animals were housed in individually ventilated Tecniplast cages at 20–24 °C and 45–65% relative humidity under a 12-h light–dark cycle, with unrestricted access to SAFE 105 diet and water.

### In vivo LPG rescue

The in vivo LPG rescue was performed as established previously (*1*) and is summarized here. *S,S*-C18:1-LPG, in which both glycerol moieties are *S*-configured, was custom-synthesized (WuXi AppTec) by the route reported in the same study (*1*); the *R,rac*-C18:1-LPG control, which cannot enter BMP synthesis in the absence of CLN6, was purchased from Avanti (858125). The lipids were dissolved in chloroform and the solvent evaporated under a stream of nitrogen. The resulting lipid films were reconstituted at 20 mM in 12% fatty acid- and endotoxin-free BSA (Sigma, A8806) prepared in 150 mM NaCl. Samples were gently warmed and sonicated to produce homogeneous lipid–BSA complexes. The vehicle consisted of the same BSA solution without lipid.

Conscious seven- to eight-week-old littermates received a single 100 µl intravenous injection via the tail vein containing 2 µmol BSA-conjugated *S,S*-C18:1-LPG, 2 µmol BSA-conjugated *R,rac*-C18:1-LPG or the corresponding BSA vehicle. Mice were randomly allocated to the following groups, with sex considered during allocation and both sexes represented in each group: wild-type mice receiving BSA vehicle (n = 7; two females and five males), *Cln6^nclf/nclf^* mice receiving BSA vehicle (n = 7; two females and five males), *Cln6^nclf/nclf^* mice receiving *R,rac*-C18:1-LPG (n = 7; five females and two males), and *Cln6^nclf/nclf^*mice receiving *S,S*-C18:1-LPG (n = 8; four females and four males). Mice were euthanized by cervical dislocation 16 h after injection, and livers were rapidly collected and snap-frozen for analysis.

### Human cell culture

HeLa cells (ECACC 93021013) and Flp-In T-REx 293 cells (Thermo Fisher Scientific, R78007) were maintained at 37 °C in humidified 5% CO2 in DMEM supplemented with 10% fetal bovine serum. Cultures were passaged with TrypLE Express at approximately 90% confluence and were routinely screened for mycoplasma.

Transient transfections were carried out in complete growth medium using FuGENE HD. For one well of a six-well plate, 0.4 µg plasmid DNA was combined with 1.2 µl FuGENE HD in 20 µl Opti-MEM. Medium was exchanged the following day, and cells were analyzed 48-72 h after transfection.

For lipid measurements, confluent six-well cultures were transferred to ice, washed twice with 3 ml ice-cold PBS and scraped into 500 µl ice-cold PBS. Cell suspensions were collected in pre-chilled 2-ml tubes and centrifuged at 500 × g for 5 min at 4 °C. After removal of PBS, pellets were snap-frozen and stored at -70 °C. Each independently processed culture well was treated as one biological replicate.

### Molecular biology and site-directed mutagenesis

CLN6 substitutions were introduced into the relevant mammalian expression plasmids using the Q5 Site-Directed Mutagenesis Kit (New England Biolabs) according to the manufacturer’s instructions. The same approach was used to generate the active-site alanine substitutions, the cysteine-accessibility constructs and the disease-associated variants. All coding regions and introduced substitutions were confirmed by Sanger sequencing (Source Genomics).

### Generation and validation of CLN6, CLN5 and CLN5/6 knockout HeLa populations

Polyclonal knockout cell populations were generated by transfecting HeLa cells with ribonucleoprotein complexes assembled from recombinant TrueCut Cas9 v2 protein and synthetic TrueGuide guide RNAs, using Lipofectamine CRISPRMAX according to the manufacturer’s protocol for HeLa cells. CLN6 was targeted with a dual-guide strategy using the guide RNAs 5′-GGUUCUCACGGACAGACAGG-3′ and 5′- GAUGAUGCUCACGUACGUGA-3′, and CLN5 was targeted with the dual-guide RNA described previously (*1*): 5′-GAAAUGUACAGUUGCCAAGU-3′ and 5′- GCCUUGAUUACACCAGAAAG-3′. Matched control populations were generated in parallel using a non-targeting guide RNA. Cells were expanded without single-cell isolation to avoid clone-specific artefacts, cryopreserved and used at low passage after thawing. Loss of CLN6 and/or CLN5 was confirmed by immunoblotting and by LC-MS quantification of the summed abundance of two CLN6-specific or four CLN5-specific peptides.

### Generation and expression of inducible Flp-In T-REx 293 cell lines

Flp-In T-REx 293 cells were maintained in DMEM with GlutaMAX, 10% fetal bovine serum and 1% penicillin–streptomycin, and were passaged at approximately 90% confluence with TrypLE Express. Stable lines expressing C-terminally FLAG-STREP-tagged wild-type CLN6, CLN8 or the disease-associated CLN6 variants were generated by co-transfecting cells in six-well plates with pOG44 and the corresponding pcDNA5/FRT/TO construct at a 9:1 mass ratio in Opti-MEM using FuGENE HD. Cells were transfected at approximately 25% confluence with 0.4 µg total DNA and 1.2 µl FuGENE HD in 20 µl Opti-MEM. Medium was changed after 24 h and cells were expanded before selection for 2 weeks in complete medium containing 50 µg/ml hygromycin B, with selection medium replaced every 2–3 days. Resistant pools were expanded and cryopreserved in 90% fetal bovine serum and 10% DMSO. Inducible expression was verified by immunoblotting paired uninduced and induced cultures.

For protein production, cells were expanded in 15-cm dishes – typically 20 dishes per construct for CLN6, 10 for CLN8 – and expression was induced with 1 µg/ml tetracycline for 48–72 h. When CLN6 mutant proteins were prepared, wild-type CLN6 was cultured, induced and harvested alongside them as a matched control. Cells were washed with ice-cold PBS, scraped into PBS on ice, pelleted, washed once more in PBS and stored at −70 °C.

### Cell treatments

For d_5_-PG tracing, 16:0-18:1 d_5_-PG (Avanti Polar Lipids, cat. no. 860385) was prepared as a 10 mM stock in ethanol. Immediately before treatment, the stock was added dropwise to pre-warmed complete DMEM and mixed thoroughly to obtain a final concentration of 25 µM. Cells were labeled for 4 h before collection for lipid analysis.

For GPG rescue experiments, *rac,rac*-[^13^C_5_]GPG was synthesized as described previously (*2*), whereas unlabeled *R,rac*-GPG was prepared as separately described below. Cells were treated with 250 µM *rac,rac*-[^13^C_5_]GPG, which contains the *S,S*-[^13^C_5_]GPG stereoisomer, or with 250 µM unlabeled *R,rac*-GPG. Cells were collected for lipid analysis after 18 or 72 h. For the 72-h GPG treatment, medium containing fresh GPG was replaced every other day. For LPG rescue experiments, *S,S*-C18:1-LPG was synthesized as described previously (*1*), and *R,rac*-C18:1-LPG was purchased from Avanti (858125). Lipids were reconstituted in BSA as described above, except that 10 mM lipid-BSA stocks were prepared, and were added to the culture medium at a final concentration of 10 µM. Cells used for lipid measurements were collected after 16 h. For whole-cell proteomic analysis, CLN6 KO cells were treated separately with each LPG stereoisomer for 48 h, with matched BSA-treated control and CLN6 KO cells analyzed in parallel.

### Substituted-cysteine accessibility mapping of CLN6 topology

Substituted-cysteine accessibility mapping (SCAM) was performed in six-well plates. HeLa cells were seeded and transfected with the desired constructs at 15–20% confluence, and the assay was performed 48 h post-transfection. Streptolysin O (SLO) was activated with 10 mM Tris(2-carboxyethyl)phosphine hydrochloride (TCEP) for 30 min at 37 °C prior to use. Cells were washed with ice-cold SHPM buffer (250 mM sucrose, 20 mM HEPES, 110 mM potassium acetate, 2.5 mM magnesium chloride, pH 7.4), then incubated with 1 ml of ice-cold SLO working solution (800 U/ml) for 10 min to allow membrane binding. Unbound SLO was removed by two washes with 2 ml ice-cold SHPM buffer. Pore formation was triggered by adding 1 ml of pre-warmed (37 °C) SHPM buffer and transferring the plate immediately to a 37 °C incubator for 10 min. Residual SLO and TCEP were removed by four washes with 2 ml ice-cold SHPM buffer. Unpermeabilised controls were processed in parallel without SLO, and fully permeabilized controls were treated with 0.1% Triton X-100 in PBS for 10 min at room temperature in place of SLO treatment.

Permeabilized cells were labeled by adding 1 ml of 100 µM MAL-dPEG₁₂-Tris(-dPEG₂₄- acid)₃ maleimide reagent (diluted from stock in SHPM) per well and incubating for 60 min at room temperature on a rocker, with the plate wrapped in foil to protect from light. Unreacted maleimide reagent was quenched by a single wash with 2 ml ice-cold SHPM, followed by addition of 1 ml of 2.5 mM L-cysteine in SHPM for 10 min on ice; residual reagents were removed by two further washes with 2 ml ice-cold SHPM.

Cells were lysed immediately thereafter in RIPA buffer supplemented with 5 mM DTT and protease inhibitor cocktail (100 µl per well). Cells were scraped and collected in 1.5 mL tubes, then kept on ice for 10 min to allow lysis. Lysates were clarified by centrifugation at 1,000 × g for 5 min at 4°C, and supernatants were collected for downstream immunoblotting.

### Purification of CLN6 and CLN8

Thawed cell pellets were resuspended in 20 mM HEPES, pH 7.4, containing 150 mM NaCl, and cellular membranes were solubilized for 2 h at 4 °C by adding lauryl maltose neopentyl glycol (LMNG) to a final concentration of 1.5%. Insoluble material was removed by centrifugation at 128,000 × g for 1 h at 4 °C. Strep-Tactin XT 4Flow resin (600 µl slurry) was washed with water and equilibrated in 20 mM HEPES, pH 7.4, 150 mM NaCl and 0.1% LMNG. The clarified extract was incubated with the resin for 2 h at 4 °C, after which the resin was washed three times with the same buffer. Bound protein was eluted in 1 ml of 50 mM HEPES, pH 7.4, 150 mM NaCl, 0.05% LMNG and 100 mM biotin. Free biotin was removed using a PD MiniTrap G-25 column, protein concentration was determined by absorbance at 280 nm, and aliquots were stored at −70 °C. Wild-type CLN6, the alanine-scan and cysteine-insertion constructs, the disease-associated CLN6 variants and CLN8 were all purified by this procedure, with wild-type CLN6 purified in parallel as a matched control whenever mutant proteins were prepared.

### Preparation of GPG stereoisomers

*R,S*-GPG for the in vitro assays was custom-synthesized (o2h Group) as described previously (*1*). *rac,rac*-[^13^C_5_]GPG, a stereoisomeric mixture carrying a uniformly ^13^C_5_-labeled glycerol headgroup and containing the *S,S*-[^13^C_5_]GPG stereoisomer, was synthesized as described previously (*2*).

*R,R*-GPG, *S,S*-GPG, *R,rac*-GPG and the d_5_-*R,rac*-GPG tracer were each prepared by mild alkaline methanolysis of a corresponding acylated lipid, following the procedure described previously for BMP (*1*). The starting lipids were, respectively, *R,R*-BMP (Avanti, 857136), *S,S*-BMP (Avanti, 857135), *R,rac*-C18:1-LPG (Avanti, 858125) and 17:0 lyso-PG-d_5_ (Avanti, 858130L). In each case, 5 mg lipid was dissolved in 1 ml of 0.1 M NaOH in methanol and incubated for 1 h at 37 °C, and the reaction was neutralized with 1 M acetic acid. Fatty acid methyl esters and residual acylated lipid were removed by butanol–methanol partitioning, and the GPG-containing aqueous phase was collected and dried under nitrogen. The residue was redissolved in methanol, insoluble salts were removed by paper filtration, and the filtrate was dried again under nitrogen. Each preparation was dissolved in water, aliquoted and stored at −20 °C. Concentrations were determined by LC-MS/MS against the chemically synthesized *R,S*-GPG standard, and the absence of residual BMP or LPG was confirmed by LC-MS/MS. In the d_5_ tracer, the deuterium label is carried on the *R*-configured glycerol moiety.

### Two-step CLN6–CLN8 stereoconversion assay

CLN6-dependent conversion of *R,S*-GPG to *S,S*-GPG was measured in a two-step assay in which purified CLN8 selectively acylates the *S,S*-GPG product to form C22:6-LPG (*1*). Reactions were assembled on ice in 1.5-ml microcentrifuge tubes. Unless otherwise indicated, the first step contained 2 µg CLN6 and 100 µM synthetic *R,S*-GPG in 25 µl of reaction buffer comprising 20 mM HEPES, pH 7.4, 150 mM NaCl, 0.05% LMNG and 5 mM CaCl2. Samples were incubated at 37 °C for 90 min, or for the indicated duration in time-course experiments. CLN6 was then heat-inactivated at 70 °C for 10 min, and samples were transferred to ice.

The second step was initiated by adding 3.5 µl of a mixture containing 100 ng CLN8 and 100 µM docosahexaenoyl-CoA (C22:6-CoA), giving a total reaction volume of 28.5 µl. After incubation at 37 °C for 15 min, reactions were quenched with 200 µl of 80% acetonitrile containing 0.2% formic acid. C22:6-LPG formation was quantified by LC-MS/MS. To characterize the reaction, the first step was performed with the indicated CLN6 amounts, incubation times, Ca^2+^ concentrations or GPG stereoisomers (*R,S*, *S,S* or *R,R*). Alternative metal ions were tested at 5 mM. Isotope-tracing experiments used d_5_-*R,rac*-GPG in the presence or absence of CLN6; diagnostic LC-MS/MS fragments identified whether C22:6 was attached to the labeled or unlabeled glycerol moiety.

Disease-associated CLN6 variants were compared with wild-type CLN6 and the H178A control using a modified first step containing 200 µM *R,S*-GPG in 21.5 µl. These reactions were incubated for 30 min, within the initial linear phase of the time course. Addition of the 3.5-µl CLN8 mixture gave a final volume of 25 µl. All other assay conditions were as described above.

Reactions lacking CLN8 served as controls for the acylation step, whereas reactions containing CLN8 but no CLN6 defined the background signal. Relative CLN6 activity was calculated as 100 × (variant LPG signal − no-CLN6 signal)/(wild-type LPG signal − no-CLN6 signal). Negative values therefore indicate product signals below the CLN8-only background. Each data point represents a separate enzymatic reaction.

### Calcium Green-5N competition assay

GPG binding to Ca^2+^ was assessed by competition with the low-affinity fluorescent indicator Calcium Green-5N. Calcium Green-5N (1 µM) was equilibrated with 10 µM CaCl_2_ in 3 ml of 1 mM NH₄HCO₃, pH 7.8, in a quartz cuvette. Fluorescence was measured on a Shimadzu RF-5301PC fluorimeter using excitation at 506 ± 1.5 nm and emission detection at 532 ± 1.5 nm. Two sequential additions of water or 50 µM *R,S*-GPG, glycerophosphocholine or glycerophosphoethanolamine were made at the times indicated in the fluorescence traces.

### Protein extraction, SDS-PAGE, Coomassie staining and immunoblotting

For immunoblotting, cells in six-well plates were placed on ice, the medium was removed, and each well was washed with 2 ml ice-cold PBS. Cells were scraped into 100 µl RIPA buffer supplemented with protease inhibitors and incubated on ice for 10 min. Lysates were clarified at 1,000 × g for 5 min at 4 °C, and the supernatants were retained.

Pre-weighed frozen tissue was homogenized with 1.4-mm ceramic beads (Cole-Parmer, 19-645-3) in 0.5 ml ice-cold RIPA buffer containing protease inhibitors. Homogenates were incubated on ice for 30 min and clarified at 5,000 × g for 15 min at 4 °C. Protein concentrations in cell and tissue lysates were measured using the Pierce BCA Protein Assay, except for SCAM samples, which were measured using the Bio-Rad Quick Start Bradford Protein Assay.

Protein purification fractions, cellular and liver lysates were resolved on 12% SDS-polyacrylamide gels cast with SureCast 40% acrylamide solution. Samples were mixed at room temperature with Lane Marker Reducing Sample Buffer to a final concentration of 2.5× and were not heated, because heating promotes aggregation of multi-pass membrane proteins. Electrophoresis was performed in Tris-glycine-SDS buffer at 120 V through the stacking gel and 180 V through the resolving gel.

For Coomassie staining, gels were immersed overnight in 15 ml InstantBlue and destained in water before scanning. For immunoblotting, proteins were transferred to 0.45-µm PVDF in chilled Tris-glycine buffer containing 10% methanol at 90 V for 100 min using a Mini Trans-Blot apparatus with an ice pack. Membranes were blocked in 5% skimmed milk in TBST, incubated with primary antibody (1:10,000 dilution for anti-vinculin and anti-GAPDH; 1:1,000 for all other antibodies) overnight at 4 °C, washed and then incubated for 1 h at room temperature with the appropriate secondary antibody diluted 1:5,000. Enhanced chemiluminescence was detected with a ChemiDoc MP system. Where required, chemiluminescent and colorimetric images were overlaid to show the molecular-mass marker.

### Immunofluorescence and confocal microscopy

Cells were transfected with the CLN6-HA plasmid using FuGENE HD (Promega, E2311) in Opti-MEM reduced-serum medium (Gibco, 31985062) according to the manufacturer’s instructions and were processed at least 24 h after transfection. Immunofluorescence was carried out as previously described (*5*). Briefly, cells grown on coverslips were fixed in 5% paraformaldehyde in PBS for 15 min at 37 °C, washed three times in PBS, and quenched in 50 mM ammonium chloride in PBS for 10 min. Following three washes in PBS, cells were permeabilized in 0.1% Triton X-100 (Sigma-Aldrich, T8787) in PBS for 10 min and washed three times in PBS. Cells were blocked in 10% fetal bovine serum in PBS for 20 min. Primary antibodies were applied at 1:500 in 5% fetal bovine serum in PBS for 2 h at room temperature, followed by three washes in the same buffer, and cells were incubated with secondary antibodies at 1:1,000 in 5% fetal bovine serum in PBS for 1 h at room temperature. Coverslips were washed three times in PBS, followed by a wash in deionized water, and mounted in Dako fluorescence mounting medium (Agilent Dako, S3023).

Images were acquired with a 100×, numerical aperture 1.4 objective on a Nikon Eclipse Ti-E inverted microscope fitted with an Andor Dragonfly 500 spinning-disk confocal system and a Zyla 4.2 PLUS sCMOS camera (Andor), controlled through Fusion software (Andor). Acquisition settings were held constant within each experiment to allow to allow comparison between samples. Z-stacks of seven planes separated by 0.2 µm were acquired and combined by maximum-intensity projection in Fiji, and the “Smooth” function was applied once to all displayed images. Line profiles were drawn through the regions indicated in the figures and used to compare the fluorescence intensities of CLN6-HA with those of calnexin, lamin B1, or GM130.

Primary antibodies for immunofluorescence were rabbit monoclonal anti-HA (clone C29F4; Cell Signaling Technology, 3724), rat monoclonal anti-calnexin (clone W17077C; BioLegend, 699401), mouse monoclonal anti-GM130 (BD Transduction Laboratories, 610822) and mouse monoclonal anti-lamin B1 (clone 119D5-F1; Invitrogen, MA1-06103), each used at 1:500. Secondary antibodies, each used at 1:1,000, were F(ab′)₂-goat anti-rabbit IgG (H+L) cross-adsorbed Alexa Fluor 488 (Invitrogen, A-11070), goat anti-mouse IgG1 cross-adsorbed Alexa Fluor 568 (Invitrogen, A-21124) and goat anti-rat IgG (H+L) cross-adsorbed Alexa Fluor 647 (Invitrogen, A-21247).

### Lipid extraction

Lipids were extracted using the butanol-methanol (BUME) procedure (*6*). Extractions were carried out in 2-ml screw-cap polypropylene tubes, with a blank tube processed alongside each batch to monitor plastic-derived background. Frozen cell pellets, lysosome-bound beads, or pre-weighed frozen tissue were homogenized in 500 µl ice-cold butanol:methanol (3:1, v/v). Tissue was disrupted with 1.4-mm ceramic beads. For untargeted lipidomic analysis, EquiSPLASH LipidoMIX was added as an internal standard before extraction, and process blanks were prepared and analysed in parallel.

After homogenization, 500 µl 1% acetic acid and 500 µl heptane:ethyl acetate (3:1, v/v) were added. Samples were vortexed for 5 min and centrifuged at 6000 × g for 5 min. The upper organic phase was transferred to a glass vial, the aqueous phase was extracted a second time and the organic phases were combined. Solvent was removed under nitrogen, and dried extracts were stored at -70 °C.

### Release and biological derivatization of GPG headgroups from liver lipids

Lipids were extracted from approximately 50 mg of liver from wild-type and Cln6nclf/nclf mice as described above. To release glycerophosphodiester headgroups, including GPG, the lipid extracts were subjected to mild alkaline methanolysis in 1 ml of 0.1 M NaOH in methanol for 1 h at 37 °C. Non-polar reaction products and residual acylated lipids were removed by butanol–methanol partitioning. The recovered aqueous fraction was dried and redissolved in water, and its GPG content was quantified by LC–MS/MS. Samples were normalized to contain equal amounts of GPG before use as substrates in a subsequent in vitro CLN8 reaction.

### Untargeted LC-MS lipidomics

Dried extracts were reconstituted in 100 µl isopropanol:acetonitrile:water (2:1:1, v/v/v) and vortexed thoroughly. A 10-µl aliquot was injected onto an Acquity Premier CSH column (2.1 × 50 mm, 1.7 µm; Waters) held at 55 °C on a Waters H-Class ACQUITY UPLC system. Mobile phase A was acetonitrile:water (60:40, v/v) containing 10 mM ammonium formate, and mobile phase B was isopropanol:acetonitrile (90:10, v/v) containing 10 mM ammonium formate. The flow rate was 500 µl/min. Mobile phase B was maintained at 40% for the first 1.5 min, increased linearly to 99% between 1.5 and 8 min, and held at 99% until 10 min. It was then returned to 40% by 10.1 min, and the column was re-equilibrated at 40% B until 12 min.

Full-scan data were acquired on a Q Exactive Orbitrap mass spectrometer equipped with heated electrospray ionization. Source settings were 300 °C capillary temperature, 420 °C heater temperature, sheath/auxiliary/sweep gas values of 40/15/3 arbitrary units and 4 kV spray voltage. Spectra were collected from m/z 120 to 1,800 at four scans per second and 35,000 resolution at m/z 200, with continuous switching between positive and negative modes.

Peaks were assigned and integrated in Xcalibur. PC, PE, PS, LPC, ePC and ePE were quantified as [M+H]^+^ ions, whereas PG, BMP and CL were quantified as [M−H]^−^ ions. Peak areas were normalized to the corresponding class-specific EquiSPLASH LIPIDOMIX internal standard; 15:0-18:1(d_7_)-PG was used for BMP and CL. For human LysoIP samples, BMP abundance was expressed relative to the combined molar abundance of all quantified PC and PE species to account for variation in lysosome input. For the HeLa phospholipidome comparison, individual phospholipid species were normalized to the corresponding internal standards and to cell-pellet protein content before differential-abundance analysis. For the volcano plot, control and CLN6 KO cells were compared using multiple unpaired t tests, and q values were calculated using the Benjamini–Krieger–Yekutieli false-discovery-rate procedure.

### GPG extraction and quantification

GPG was extracted from pre-weighed tissue in 1 ml 80% methanol using 1.4-mm ceramic beads. Homogenates were clarified at 20,000 × g for 15 min at 4 °C, and supernatants were transferred to glass vials.

GPG was measured on a Shimadzu 8060 triple-quadrupole mass spectrometer coupled to an LC-30AD system. Separation used an Atlantis Premier BEH Z-HILIC column (2.1 × 150 mm, 1.7 µm; Waters), mobile phase A consisting of 15 mM ammonium acetate with 0.1% ammonia and mobile phase B consisting of methanol. The run was performed at 0.1 ml/min with injection at 0.5 min and the following 6-min program: 80% B from 0 to 1 min, 80% to 20% B from 1 to 3 min, 20% B from 3 to 5 min and 80% B from 5 to 6 min. Source settings were nebulizing gas 3 l/min, heating gas 10 l/min, interface temperature 300 °C, desolvation-line temperature 245 °C, heat block temperature 400 °C and drying gas 10 l/min. Negative-ion multiple-reaction monitoring used m/z 244.5→152.9 (100-ms dwell, Q1 20 V, collision energy 15 eV, Q3 20 V), 244.5→78.95 (100-ms dwell, Q1 20 V, collision energy 35 eV, Q3 20 V) and 244.5→170.95 (100-ms dwell, Q1 20 V, collision energy 17 eV, Q3 21 V). Concentrations were calculated against the synthetic *R,S*-GPG standard.

### Targeted LC-MS/MS analysis of lipids

Targeted lipid analysis followed the previously published LC-MS/MS workflow (*1*), with the chromatography and source conditions given below. The established positive-ion transitions, cone voltages and collision energies are listed in the table below.

#### Positive-ion MRM transitions used for targeted lipid analysis

| Analyte | Precursor $m/z$ | Product $m/z$ | Cone voltage (V) | Collision energy (eV) |
| --- | --- | --- | --- | --- |
| C14:0-LPG | 457.4 | 285.3 | 50 | 20 |
| C16:0-LPG | 485.4 | 313.3 | 50 | 20 |
| C18:2-LPG | 509.4 | 337.3 | 50 | 20 |
| C18:1-LPG | 511.4 | 339.3 | 50 | 20 |
| d<sub>5</sub>-C18:1-LPG (headgroup) | 516.4 | 339.3 | 50 | 20 |
| d<sub>5</sub>-C18:1-LPG (backbone) | 516.4 | 344.3 | 50 | 20 |
| C18:0-LPG | 513.4 | 341.3 | 50 | 20 |
| C20:4-LPG | 533.4 | 361.3 | 50 | 20 |
| C22:6-LPG | 557.5 | 385.3 | 50 | 20 |
| d<sub>5</sub>-C22:6-LPG (headgroup) | 562.5 | 385.3 | 50 | 20 |
| d<sub>5</sub>-C22:6-LPG (backbone) | 562.5 | 390.3 | 50 | 20 |
| 16:0-18:1-BMP | 749.6 | 313.3, 339.3 | 50 | 20 |
| 18:2-18:1-BMP | 773.6 | 337.3, 339.3 | 50 | 20 |
| 18:2-18:2-BMP | 771.6 | 337.3 | 50 | 20 |
| 18:1-18:1-BMP [M+H]<sup>+</sup> | 775.6 | 339.3 | 50 | 20 |
| 18:1-18:1-BMP [M+Na]<sup>+</sup> | 797.6 | 459.3 | 50 | 20 |
| 18:1-18:0-BMP | 777.6 | 339.3, 341.3 | 50 | 20 |
| 16:0-22:6-BMP | 795.6 | 313.3, 385.3 | 50 | 20 |
| 18:1-20:4-BMP | 797.6 | 339.3, 361.3 | 50 | 20 |
| 18:2-22:6-BMP | 819.6 | 337.3, 385.3 | 50 | 20 |
| 18:1-22:6-BMP [M+H]<sup>+</sup> | 821.6 | 339.3, 385.3 | 50 | 20 |
| 18:1-22:6-BMP [M+Na]<sup>+</sup> | 843.6 | 459.3, 505.3 | 50 | 20 |
| 20:4-22:6-BMP | 843.6 | 361.3, 385.3 | 50 | 20 |
| 22:6-22:6-BMP [M+H]<sup>+</sup> | 867.7 | 385.3 | 50 | 20 |
| 22:6-22:6-BMP [M+Na]<sup>+</sup> | 889.7 | 505.3 | 50 | 20 |

Multiple product ions listed in one cell denote separate transitions. “Headgroup” and “backbone” identify the location of the d_5_ label in the non-acylated and acylated glycerol moieties, respectively.

LPG and BMP were quantified on a Xevo TQ-S triple-quadrupole mass spectrometer coupled to an I-Class ACQUITY UPLC. Samples were held in glass vials at 8 °C, and 10 µl was injected onto a BEH C18 column (1 × 50 mm, 1.7 µm, 130 Å; Waters) held at 30 °C and protected by a 0.2-µm in-line filter. Mobile phase A was 5% acetonitrile with 0.1% formic acid, and mobile phase B was 95% acetonitrile with 0.1% formic acid. The flow rate was 0.2 ml/min. Mobile phase B was held at 5% for 1.5 min, increased linearly to 40% at 3 min, 70% at 12 min and 100% at 15 min, and then maintained at 100% for the remainder of the run. The total run time was 20 min for LPG-only analyses and 60 min when both LPG and BMP were measured.

Analytes were detected in positive-ion electrospray mode by multiple-reaction monitoring. The capillary voltage was 3.1 kV, source temperature 150 °C and desolvation temperature 500 °C; nitrogen and argon were used as curtain and collision gases, respectively. Peaks were integrated in MassLynx 4.1 using automated peak selection followed by manual review. LPG and BMP were quantified using five-point, ten-fold serial calibration curves prepared from *R,rac*-LPG (Avanti 858125) and *R,S*-BMP (Avanti 857133), respectively. Multiple transitions with matching retention times were used to validate peak identity. For BMP, the [M+Na]^+^ transition was used where its greater sensitivity was advantageous.

The assay did not resolve BMP stereoisomers or acyl positional isomers; structural drawings therefore show the expected biological configuration rather than a chromatographically assigned configuration. *S,S*-C18:1-LPG and *R,rac*-C18:1-LPG were also not distinguished by this LC-MS/MS workflow. In contrast, LPG 1-acyl and 2-acyl positional isomers eluted as separate peaks, with the 2-acyl species eluting first (*1*). Only the first LPG peak was produced in recombinant CLN8 assays and was used for quantification. For biological samples, both LPG positional-isomer peaks were summed where relevant. For d_5_-C22:6-LPG, the transitions *m/z* 562.5→385.3 and 562.5→390.3 distinguished isotope labelling in the non-acylated glycerol headgroup and the acylated glycerol backbone, respectively (see the transition table above).

For BMP species carrying different acyl chains on the two glycerol moieties, collision-induced dissociation of a single precursor yields two monoacylglycerol product ions, formed by cleavage of the phosphodiester bond on either side of the phosphate. For 18:1-22:6-BMP ([M+H]^+^, precursor *m/z* 821.6), cleavage between the phosphate and the 22:6- or 18:1- bearing glycerol produces ions at *m/z* 385.3 and 339.3, respectively (see transition table above). Both transitions were acquired in the same run, and their relative yields were expressed as the ratio of their peak areas (385.3/339.3). For the in vivo LPG rescue experiment, the corresponding sodium-adduct precursor ([M+Na]⁺, *m/z* 843.6) was used, and the product-ion peak-area ratio was calculated as 505.3/459.3. These product ions retain the sodiated phosphate group: cleavage between the phosphate and the 22:6-bearing glycerol generates the ion at *m/z* 459.3, whereas cleavage between the phosphate and the 18:1-bearing glycerol generates the ion at *m/z* 505.3 (see transition table above).

### Quantitative proteomics of human cells

For each replicate, cells harvested from one 10-cm plate were pelleted and lysed in 100 µl of 1% sodium deoxycholate, 100 mM triethylammonium bicarbonate, 10% isopropanol and 50 mM NaCl containing protease/phosphatase inhibitors and nuclease. Lysates were briefly sonicated, and protein was measured by Bradford assay. Ten micrograms of protein were reduced and alkylated for 1 h at room temperature in the dark with 5 mM tris(2-carboxyethyl)phosphine and 10 mM iodoacetamide, followed by overnight digestion at room temperature with 500 ng trypsin. Sodium deoxycholate was precipitated by acidification with trifluoroacetic acid and removed at 12,000 × g. Peptides were desalted on C18 spin tips, dried and reconstituted in 40 µl 0.1% formic acid; approximately 500 ng was injected.

Peptides were separated on a Vanquish Neo UHPLC and analyzed with an Orbitrap Fusion Lumos. After trapping on a 300 µm × 5 mm C18 cartridge, peptides were resolved over 100 min on a 75 µm × 50 cm C18 column at 300 nl/min. The same sample-preparation and acquisition workflow was used for both the control-versus-CLN6 KO comparison and the LPG-rescue experiment. Full MS scans were acquired from *m/z* 375 to 1,200 at 60,000 resolution, followed by data-independent acquisition at 30,000 resolution using 42 windows spanning *m/z* 400-900 and HCD collision energy 30%.

Raw data were processed in DIA-NN 2.0 against an in-silico-predicted human spectral library generated from FASTA release 9606_2025_01_23. Searches allowed one missed cleavage, N-terminal methionine excision and fixed cysteine carbamidomethylation. Peptide length was limited to 7-30 residues, precursor charge to 1-4, precursor *m/z* to 375-1,200 and fragment *m/z* to 145-1,450. Match-between-runs and retention-time-dependent cross-run normalization were enabled, using QuantUMS high-precision quantification. For knockout validation, the summed abundance of two unique CLN6 peptides or four unique CLN5 peptides was used.

### Quantitative proteomics of mouse livers

Frozen mouse liver tissue was homogenized in ice-cold RIPA buffer, and lysates were clarified by centrifugation at 20,000 × g for 10 min. Protein concentrations in the supernatants were determined using a BCA assay. Ten micrograms of liver protein in 40 µl were reduced and alkylated with 10 mM tris(2-carboxyethyl)phosphine and 20 mM 2-chloroacetamide for 10 min at 80 °C, followed by 15 min at room temperature. Proteins were processed by single-pot, solid-phase-enhanced sample preparation (SP3) using a KingFisher Apex equipped with a 96-deep-well magnetic head. Methanol (160 µl) containing 100 µg SP3 beads was added to each sample. Beads consisted of a 1:1 mixture of Cytiva 45152105050250 and 65152105050250, washed three times in LC-MS-grade water before use. Samples were mixed using the instrument’s slow setting for 5 min. Beads were collected and washed three times, without release, in 1 ml 80% ethanol. Proteins were digested and peptides released into 150 µl of 20 mM ammonium bicarbonate containing 400 ng trypsin/Lys-C (Promega, V5073) for 2 h at 47 °C. At each collection step, beads were gathered three times for 10 s. Peptide solutions were acidified with 10 µl 1% trifluoroacetic acid, incubated for 10 min at 4 °C and clarified at 500 × g for 5 min.

Peptides equivalent to 500 ng pre-digestion protein were injected using a Vanquish Neo HPLC onto a PepMap Neo C18 trapping cartridge (300 µm × 5 mm). Separation used a PepMap RSLC C18 EasySpray column (150 µm × 15 cm, 2 µm, 100 Å) at 50 °C and 1.8 µl/min. Mobile phase A was 0.1% formic acid, and mobile phase B was 80% acetonitrile containing 0.1% formic acid. The 11.4-min gradient increased B from 3% to 35% over 0–10.4 min and from 35% to 55% over 10.4–11.4 min.

Peptides were analyzed on an Orbitrap Astral mass spectrometer with electrospray ionization at 1.9 kV, an RF lens setting of 40% and an ion-transfer-tube temperature of 290 °C. Orbitrap MS scans covered m/z 380–980 at 240,000 resolution, with an automatic gain control (AGC) target of 5 × 106 and a maximum injection time of 5 ms. FAIMS Pro was operated at a compensation voltage of −48 V. Astral data-independent acquisition used optimized 3-Th windows, 25% HCD collision energy, an AGC target of 5 × 104 and a maximum injection time of 5 ms, with fragment acquisition over m/z 150–2,000.

Raw data were processed in DIA-NN 2.5 using a spectral library built from the Mus musculus UniProt proteome (release 2026_01) and the full universal contaminants list (PMID: 35793413). Searches allowed one missed trypsin/P cleavage, N-terminal methionine excision as one variable modification, and fixed cysteine carbamidomethylation. Peptide length was restricted to 7–30 residues, precursor charge to 1–4, precursor m/z to 380–980 and fragment *m/z* to 150–2,000. MS1 and MS2 mass-accuracy settings were 4 and 10, respectively. Protein-level false discovery rate was set to 1%, match-between-runs was enabled, and QuantUMS precision quantification was used; other settings were left at their defaults.

### Proteomic data analysis

Each proteomic dataset was analyzed separately. Valid protein intensities were log_2_-transformed and median-normalized across retained samples. Missing values were not imputed.

Mouse liver samples comprised four groups: BSA-treated wild-type mice (n = 7), BSA-treated *Cln6^nclf/nclf^* mice (n = 7), and *Cln6^nclf/nclf^* mice treated with *R,rac*-LPG (n = 7) or *S,S*-LPG (n = 8). Sample quality was assessed using protein detection rates, pairwise Pearson correlations calculated from proteins quantified in both samples, correlations with the corresponding group mean abundance profile, and principal component analysis (PCA) of proteins quantified in all samples. No outliers were identified, and all samples were retained. Because sex composition differed between groups and PCA revealed substantial sex-associated variation, all differential-abundance analyses included sex as a covariate. For each liver protein, differential abundance was assessed by ordinary least-squares linear regression of normalized log₂ intensities. The comparison between BSA-treated wild-type and *Cln6^nclf/nclf^* mice included genotype and sex as predictors; the comparison between *R,rac*- LPG- and *S,S*-LPG-treated *Cln6^nclf/nclf^*mice included treatment and sex. Genotype and treatment coefficients therefore represented sex-adjusted log_2_ fold changes. Proteins were tested only if at least three valid measurements were available in each comparison group and the model matrix remained full rank after excluding missing observations. Coefficients were evaluated using two-sided t tests based on each model’s residual variance. For each comparison, *P* values were adjusted across all successfully tested proteins using the Benjamini–Hochberg false-discovery rate procedure.

The HeLa control-versus-CLN6 KO comparison shown in Fig. 4A was acquired as an independent experiment with six biological replicates per group. Intensities were processed as described above, and differential abundance was assessed proteome-wide, with Benjamini– Hochberg correction across all tested proteins.

The HeLa rescue experiment comprised BSA-treated control cells and CLN6 KO cells treated with BSA, *R,rac*-LPG or *S,S*-LPG, initially with three biological replicates per group. One BSA-treated CLN6 KO replicate was excluded before normalization because it had substantially fewer quantified proteins, correlated poorly with the other replicates in that group and separated markedly by PCA. The final sample sizes were therefore n = 2 for BSA-treated CLN6 KO cells and n = 3 for each remaining group. SMPD1, CTSD, PPT1 and TPP1 were prespecified as proteins of interest in the HeLa rescue experiment. The primary comparison, *S,S*-LPG versus *R,rac*-LPG treatment, was assessed using two-sided Welch’s t tests on normalized log_2_ intensities. *P* values were adjusted across these four protein comparisons using the Benjamini–Hochberg procedure, with *q* < 0.05 considered significant. Other pairwise comparisons were exploratory and were not included in this four-test correction. For visualization, normalized log_2_ intensities were back-transformed to the linear scale and expressed relative to the mean abundance of each protein in control cells. Graphs show individual biological replicates.

### Acid sphingomyelinase activity assay

Liver tissue was homogenized in Milli-Q water containing cOmplete EDTA-free protease inhibitors (Roche, 04693159001) using a gentleMACS Octo Dissociator and M-tubes (Miltenyi Biotec). Homogenates were sonicated for 10 cycles of 15 s on and 15 s off using a Bioruptor Next Gen sonicator (Hologic Diagenode), then clarified by centrifugation at 10,000 × g for 10 min at 4 °C. Supernatant protein concentrations were determined by BCA assay and adjusted to 1 µg/µl.

Acid sphingomyelinase (ASM) activity was measured using the fluorogenic substrate 6-hexadecanoylamino-4-methylumbelliferyl-phosphorylcholine (HMU-PC; Thermo Fisher Scientific, 30004056), as described previously (*7*). Each 30-µl reaction contained 10 µl liver lysate (10 µg protein) and 20 µl of 1.35 mM HMU-PC prepared in 0.1 M sodium acetate, pH 4.5, containing 0.2% (w/v) sodium taurocholate and 0.02% sodium azide, giving a final HMU-PC concentration of 900 µM. Reactions were incubated at 37 °C for 90 min and quenched with 0.2 M glycine–NaOH, pH 10.7, containing 0.2% (w/v) SDS and 0.2% (w/v) Triton X-100. Released HMU was measured using a SpectraMax iD3 plate reader at excitation and emission wavelengths of 404 and 460 nm, respectively, and quantified against a standard curve of pure HMU. Activity was expressed as pmol substrate hydrolyzed per µg protein per hour.

### AlphaFold modelling and initial GPG docking

The docking ligand was *R,S*-glycerophosphoglycerol (GPG; *sn*-glycero-3-phospho-(1′-*sn*-glycerol); CAS 185615-51-4; PubChem CID 54575349). GPG was docked into an AlphaFold 3–predicted CLN6 structure (UniProt Q9NWW5) using AutoDock Vina (version 1.2.5) (*8, 9*). Receptor and ligand coordinates were supplied in PDBQT format. The search box measured 120 × 120 × 120 Å and was centered at (x, y, z) = (0.080, 1.682, −5.560) Å. Exhaustiveness was set to 128, with a maximum of 25 output poses. The pose with the lowest Vina score (−5.8 kcal/mol) was selected for graphical representation.

### Boltz-2 co-folding of CLN6, GPG and Ca^2+^

The human CLN6 sequence (UniProt Q9NWW5; 311 residues) was co-folded as a homodimer with *R,S*-GPG (PDB Chemical Component Dictionary code CU7) and explicit Ca^2+^ ions using Boltz-2 (*10*). The number of Ca^2+^ ions was treated as a modelling variable: 25 models were generated for each of the zero-, one- and two-Ca^2+-^per-active-site conditions. Models were compared using primary Boltz metrics (PAE, PDE, pLDDT, pTM) and secondary ligand-specific interface metrics (LIS (*11*), pDockQ (*12*), pDockQ2 (*13*), ipSAE (*14*)). For pocket-specific error metrics, the pocket was defined independently in each model as all protein residues with a heavy atom within 8 Å of any GPG heavy atom. Predictions with and without binding-site specification were compared. A two-Ca^2+^ model was selected for molecular dynamics based on ligand-interface confidence and pocket error.

### Membrane-system construction and classical molecular dynamics

The CLN6 homodimer was assembled with one *R,S*-GPG and two structural Ca^2+^ ions per protomer. The two copies of the substrate were placed in opposite orientations: the *R*-glycerol faced Asp59 in chain A, whereas the *S*-glycerol faced Asp59 in chain B. His178 was assigned the HID tautomer, leaving Nε2 unprotonated and available for nucleophilic attack. Protein, lipid and ligand parameters were ff19SB (*15*), Lipid21 (*16*) and GAFF2 (*17*) with AM1-BCC (*18*) ligand charges, respectively; TIP3P (*19*) water and the Li-Merz 12-6 (*20*) ion parameters were used.

The complex was embedded with packmol/packmol-memgen (*21*, *22*) in an ER-like bilayer comprising 590 lipids (295 per leaflet): 320 POPC, 120 POPE, 60 POPS, 30 POPA and 60 cholesterol molecules. The system was solvated, followed by neutralization and adjustment to 0.15 M CaCl_2_; the four active-site Ca^2+^ ions were distinguished from bulk ions.

Molecular dynamics simulations were performed using pmemd.cuda in Amber26 (*23*). Particle-mesh Ewald electrostatics were used with a 10 Å real-space and van der Waals cut-off. Bonds involving hydrogen atoms were constrained with SHAKE, permitting a 2 fs integration step. Temperature was maintained at 303 K using Langevin dynamics, and pressure was controlled with a Berendsen barostat (2 ps relaxation) under anisotropic NPγT conditions with no applied surface tension.

The system underwent 15,000 minimization cycles with 5 kcal mol−1 Å−2 restraints on the protein backbone and structural Ca^2+^ ions, followed by 10,000 unrestrained minimization cycles. It was heated from 0 to 100 K over 50 ps in the NVT ensemble with 5 kcal mol−1 Å−2 restraints on protein, ligand and Ca^2+^ heavy atoms, then from 100 to 303 K over 200 ps under NPγT conditions with 2.5 kcal mol−1 Å−2 heavy atom restraints. This was followed by 1 ns with 2.5 kcal mol−1 Å−2 restraints on the backbone, ligand and Ca^2+^ ions and 1 ns with 1 kcal mol−1 Å−2 restraints on Cα atoms, ligand and Ca^2+^ ions. The glycerol arm in each protomer was then seated against Asp59 during a 50 ns stage using only flat-bottom distance restraints centered at 4.0 Å (50 kcal mol−1 Å−2). All restraints were removed for the 500 ns production run.

Trajectories were analyzed for simultaneous formation of two geometric features: an incoming GPG O-to-phosphorus distance of no more than 4.5 Å and a phosphorus-to-His178 Nε2 distance of no more than 4.5 Å. A frame satisfying both cut-offs was classified as a near-attack conformation. A representative chain B frame satisfying both distance criteria was selected as the starting structure for QM/MM calculations.

### QM/MM free-energy calculations

QM/MM molecular dynamics was carried out with AMBER sander v22 through the TeraChem EXTERN interface (*24*, *25*). The quantum region was described at the B3LYP-D3/6-31G(d) level, and the remainder of the system was treated with the classical parameters described above. Simulations used electronic embedding, a 1 fs time step, an 8 Å non-bonded cutoff, a Langevin thermostat at 303 K and a Berendsen barostat. SHAKE was disabled for quantum region atoms.

The quantum region comprised 96 atoms with a net charge of -3: the His178 imidazole, the complete *R,S*-GPG molecule, both structural Ca^2+^ ions, the side chains of Asp59, Asp131, Asp173, Glu174, Glu227 and Glu298, and three active-site water molecules. Cα-Cβ QM/MM boundaries were capped with link hydrogens.

The reaction was divided into two phosphoryl-transfer steps through a covalent phospho-His178 intermediate. Step 1 used the collective variable (CV1), q1 = d(P-O_leaving) - d(P- Nε2), to describe His178 attack and cleavage of the original GPG phosphoester. Step 2 used CV2, q2 = d(P-O_incoming) - d(P-Nε2), describing attack by the distal glycerol oxygen and departure of His178. The proton transfer from the incoming oxygen to the leaving oxygen occurred spontaneously during step 1.

Starting configurations for umbrella windows were generated by steered molecular dynamics (*26*). Step 1 comprised 17 windows spanning q1 = -1.0 to +1.0 Å, with 0.25 Å spacing in the basins and a finer 0.125 Å spacing across the transition state barrier; a 100 kcal mol−1 Å−2 harmonic restraint was applied to q, and the spectator P-O_incoming distance was held at 3.271 Å with a 200 kcal mol−1 Å−2 restraint. Step 2 comprised 27 uniformly spaced (0.125 Å) windows from q2 = -1.75 to +1.5 Å with a 100 kcal mol−1 Å−2 force constant and no restraint on the departing oxygen (O8). Each window was equilibrated for 0.5 ps and sampled for 5 ps. Mulliken charges were also recorded at every step.

Potential-of-mean-force profiles were reconstructed using MBAR in pymbar 4.x (*27*). Uncertainty was estimated by bootstrapping across windows. The two one-dimensional profiles were joined at their common phospho-His state. A free-energy surface for the first transfer was also constructed as a function of the individual forming- and breaking-bond distances. Bond distances and atomic charges were averaged within each window after discarding the first 1 ps of production.

First-shell coordination of both structural Ca^2+^ ions was monitored in every QM/MM umbrella window using a 3.0-Å cutoff selected from the radial distribution function. Coordinating oxygen atoms were classified as carboxylate, backbone carbonyl, GPG or water oxygens. Structural renderings were prepared in PyMOL and Blender.

### Statistics and reproducibility

No statistical method was used to determine sample size in advance. Biological replicate and animal numbers are provided in the figure legends. Unless otherwise stated, graphs show the mean ± s.e.m. together with individual observations. Two-group comparisons used unpaired, two-tailed Student’s t-tests. One-way ANOVA followed by Tukey’s or Dunnett’s multiple-comparisons test was used for experiments with more than two groups, and two-way ANOVA followed by Šídák’s test was used when two independent variables were analyzed. For the mouse rescue lipid and ASM activity analyses, treatment and sex were included as factors, with Tukey-adjusted comparisons between treatment groups. The exact test applied to each dataset is stated in its figure legend.

For purified-protein experiments, individual points represent separate assay measurements. Each principal biochemical result was reproduced with at least one independently prepared CLN6 batch; representative experiments and the number of independent purifications are identified in the legends. Concentration-response and one-phase association curves were fitted in GraphPad Prism 10.2.

Graphs and statistical analyses were prepared in GraphPad Prism 10.2. Figures were assembled in Affinity Publisher 2.4.2. Molecular visualizations were generated with PyMOL. Other computational packages and versions are specified in the corresponding modelling subsections.

---

**Movie S1. Proposed catalytic mechanism of CLN6-mediated GPG epimerization**

Animation of the modeled conversion of *R,S*-GPG to *S,S*-GPG within the CLN6 active site. The view moves from membrane-embedded CLN6 to the catalytic pocket, showing phosphate transfer through a covalent phosphohistidine intermediate involving H178. Labeled pauses indicate the reactant (R), first transition state (TS1‡), phosphohistidine intermediate (Int), second transition state (TS2‡), and product (P). GPG carbon atoms are shown in yellow, protein carbon atoms in gray, oxygen in red, nitrogen in blue, phosphorus in orange, and Ca²⁺ ions as green spheres. Selected active-site residues are labeled.

**Fig. S1.**
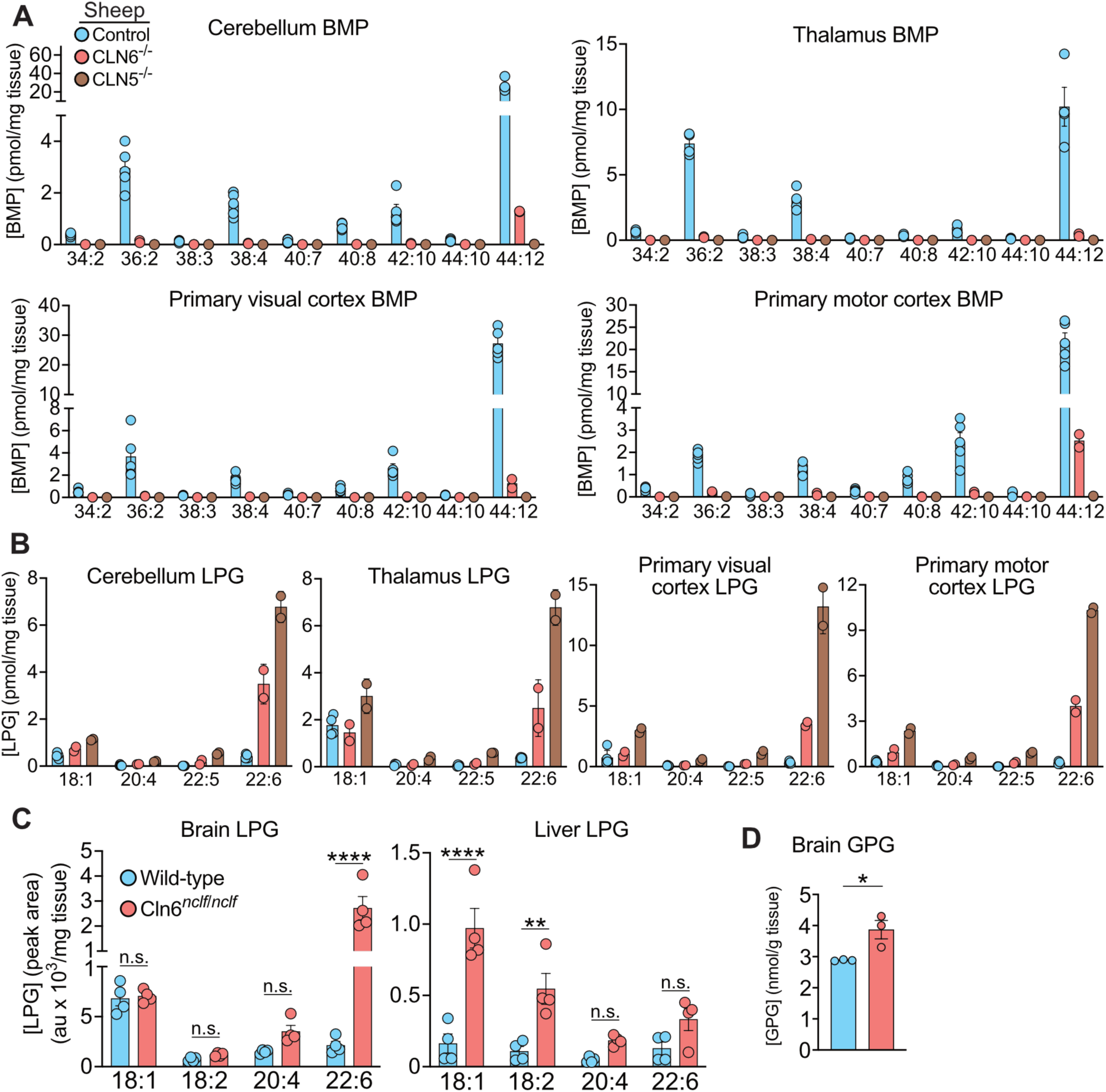
BMP and its biosynthetic intermediates in CLN6-deficient animal models. (**A** and **B**) Abundance of the indicated BMP (**A**) and LPG (**B**) species in brain regions from 12-month-old female control (n = 5), CLN6^−/−^ (n = 2) and CLN5^−/−^ (n = 2) sheep, measured by untargeted LC-MS. BMP species are labeled by the total number of acyl-chain carbons and double bonds. (**C**) Abundance of the indicated LPG species in the brain and liver of 6-month-old female wild-type and *Cln6^nclf/nclf^* mice (n = 4 per group). (**D**) Total brain GPG abundance in mice of the same age, sex and genotypes (n = 3 per group). Measurements in (**C**) and (**D**) were performed by targeted LC-MS/MS. Points represent individual animals; bars and error bars indicate means ± SEM. Statistical comparisons used two-way ANOVA with Šídák’s multiple-comparisons test in (**C**) and an unpaired two-tailed Student’s t test in (**D**). *P* < 0.05 (*), *P* < 0.01 (**) and *P* < 0.0001 (****); n.s., not significant.

**Fig. S2.**
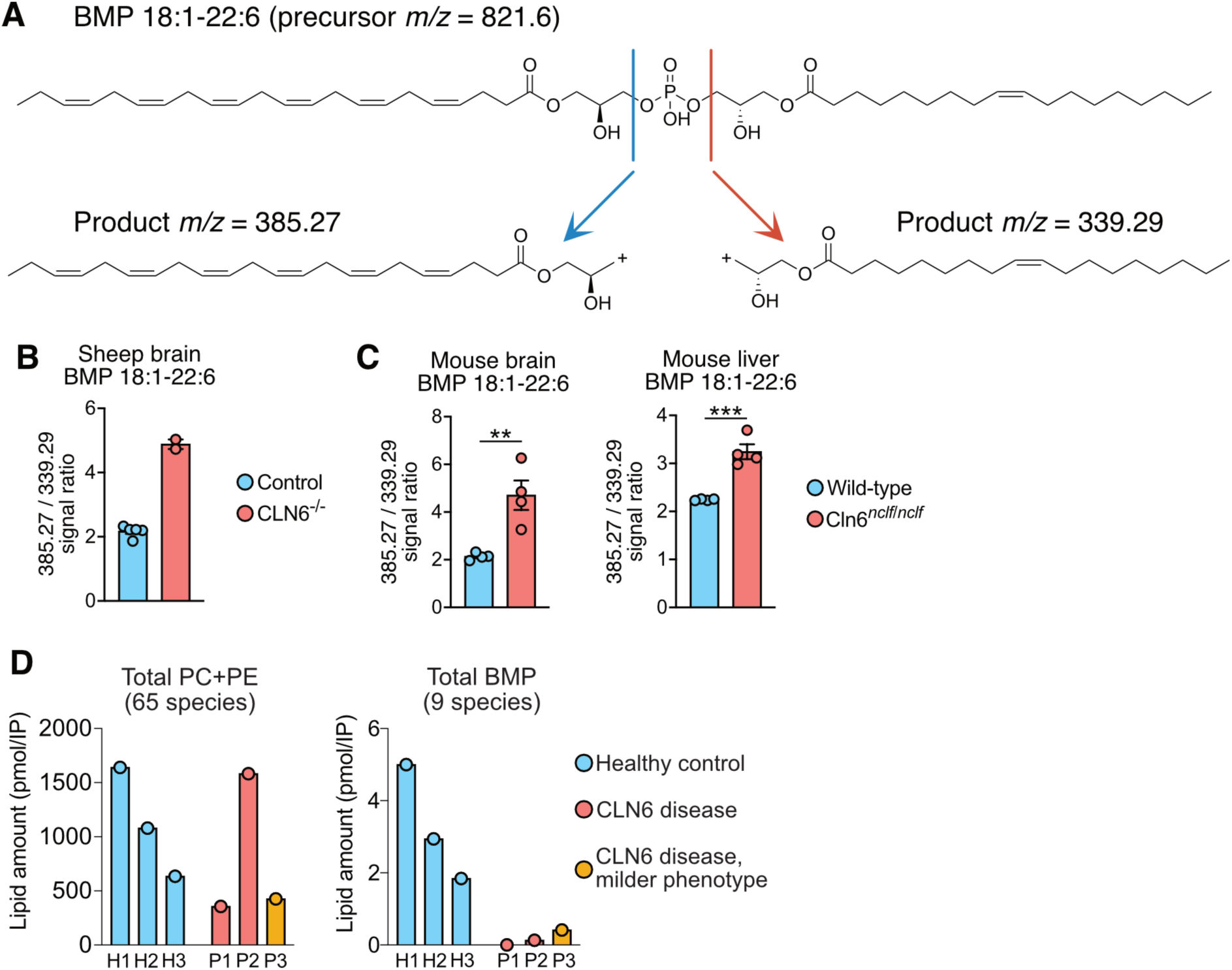
Altered BMP fragmentation in CLN6-deficient tissues and lysosomal lipid amounts in patient samples. (**A**) Fragmentation scheme for 18:1-22:6 BMP (precursor *m/z* 821.6), showing the C22:6-containing product ion at *m/z* 385.27 (blue) and the C18:1-containing product ion at *m/z* 339.29 (red). (**B** and **C**) Ratio of the signals for these two product ions (385.27/339.29), measured by targeted LC-MS/MS, in the motor cortex of control (n = 5) and CLN6^−/−^ (n = 2) sheep (**B**) and in the brain and liver of wild-type and *Cln6^nclf/nclf^* mice (n = 4 per group) (**C**); points represent individual animals; bars and error bars indicate means ± SEM. (**D**) Total amounts of phosphatidylcholine and phosphatidylethanolamine (PC + PE; 65 species combined) and BMP (9 species), measured by untargeted LC-MS in lysosomes isolated from peripheral blood mononuclear cells of three healthy controls (H1–H3) and three children with CLN6 Batten disease (P1–P3). P3, who has a milder phenotype, is highlighted in yellow. Each bar represents one individual, with lipid amounts expressed as pmol per lysosome immunoprecipitation (IP). In (**C**), *P* < 0.01 (**) and *P* < 0.001 (***) by unpaired two-tailed Student’s t tests.

**Fig. S3.**
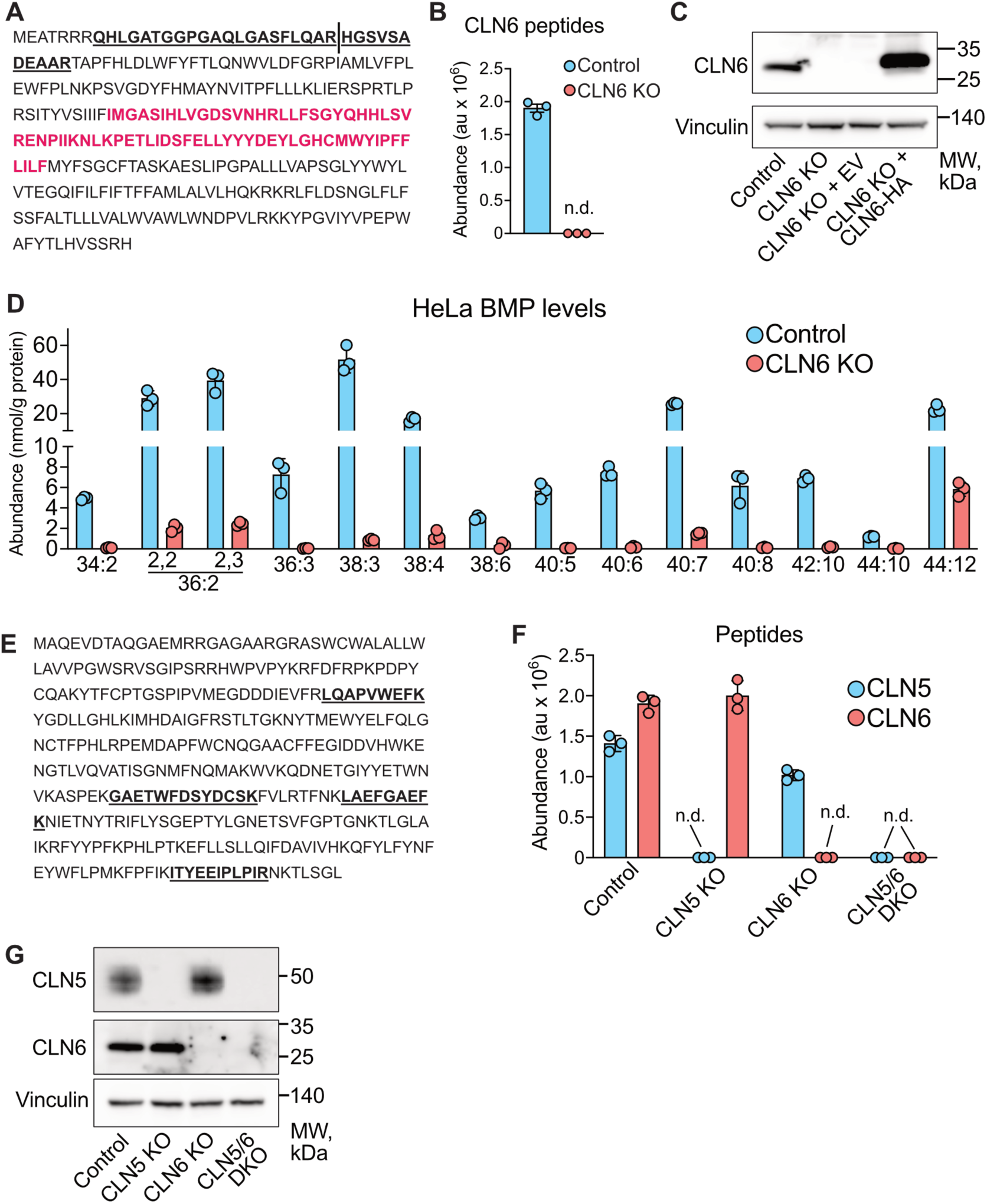
Characterization of CLN6 KO, CLN5 KO and CLN5/6 DKO HeLa cell populations. (**A**) Human CLN6 amino acid sequence, with the two peptides used for LC-MS quantification shown in bold and underlined, and the antibody immunogen sequence highlighted in red. (**B**) Summed abundance of the two CLN6 peptides in control and CLN6 KO polyclonal cell lysates (n = 3). (**C**) Immunoblot of CLN6 in control and CLN6 KO HeLa cells, including CLN6 KO cells transfected with empty vector (EV) or CLN6-HA, with vinculin as a loading control. The CLN6-HA transfection is representative of the rescue experiments in this study and illustrates transfected protein abundance relative to endogenous CLN6. (**D**) Abundance of all quantified BMP species in control and CLN6 KO HeLa cells, measured by untargeted LC-MS (n = 3). The 36:2 BMP species resolved as two neighboring peaks corresponding to the 2,2′- and 2,3′-BMP isomers. (**E**) CLN5 amino acid sequence, with the four peptides used for LC-MS quantification shown in black. (**F**) Summed abundance of the quantified peptides for CLN5 and CLN6 in control, CLN5 KO, CLN6 KO and CLN5/6 DKO polyclonal cell lysates (n = 3). (**G**) Immunoblots of endogenous CLN5 and CLN6 in the indicated cell populations, with vinculin as a loading control. In (**B**), (**D**) and (**F**), points represent individual biological replicates; bars and error bars indicate means ± SEM. n.d., not detected.

**Fig. S4.**
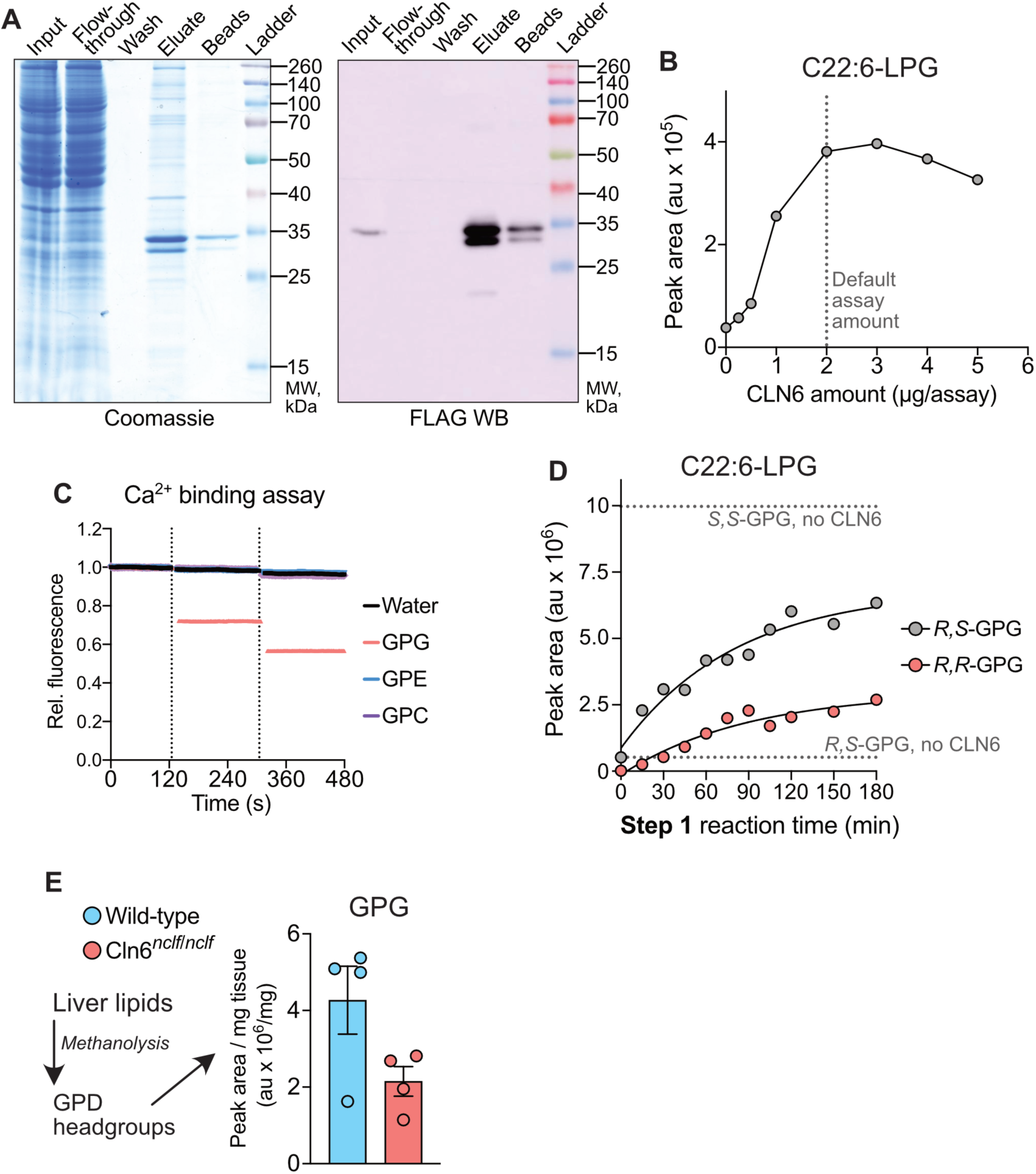
Establishment of the in vitro CLN6 stereoconversion assay. (**A**) Representative Coomassie-stained gel and anti-FLAG immunoblot of fractions collected during CLN6-FLAG-STREP purification. The eluate was used for in vitro assays. (**B**) C22:6-LPG formation using the indicated amounts of purified CLN6 in step 1 of the two-step CLN6–CLN8 assay. The dotted line marks the standard amount used in subsequent assays (2 μg). (**C**) Ca^2+^-binding assay based on competition with Calcium Green-5N. Fluorescence emission at 532 nm was monitored during two sequential additions of water or 50 μM *R,S*-glycerophosphoglycerol (GPG), glycerophosphocholine (GPC) or glycerophosphoethanolamine (GPE). Dotted lines indicate the additions; fluorescence is expressed relative to the initial signal. (**D**) C22:6-LPG formation following the indicated durations of the CLN6 reaction with *R,S*-GPG or *R,R*-GPG. Upper and lower dotted lines indicate product formation when *S,S*-GPG or *R,S*-GPG, respectively, was supplied directly to CLN8 without CLN6. Each point in (**B**) and (**D**) represents a separate enzymatic reaction. (**E**) Preparation of GPG headgroups by methanolysis of liver lipids and their abundance in samples from 6-month-old female wild-type and *Cln6^nclf/nclf^* mice (n = 4 per group), normalized to tissue mass. Points represent individual mice; bars and error bars indicate means ± SEM. C22:6-LPG in (**B**) and (**D**) and GPG in (**E**) were measured by targeted LC-MS/MS. In vitro assay results are representative of three independent CLN6 protein purifications.

**Fig. S5.**
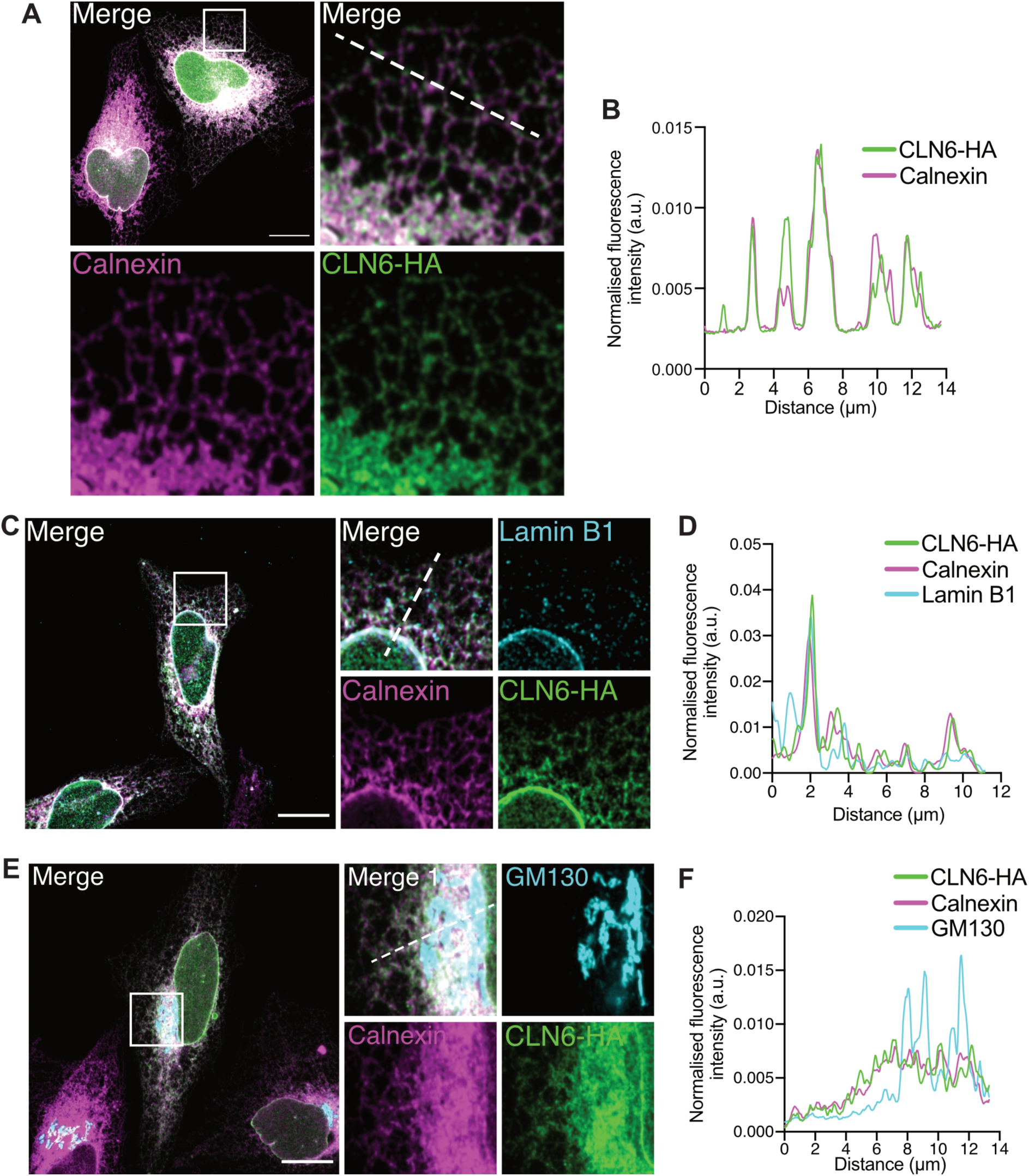
Subcellular localization of HA-tagged CLN6. (**A**, **C** and **E**) Representative confocal immunofluorescence images of HeLa cells expressing CLN6-HA, stained for HA (green) and the ER marker calnexin (magenta), together with the nuclear lamina marker lamin B1 (**C**, cyan) or the Golgi marker GM130 (**E**, cyan). Regions of interest (15 x 15 μm) in the overview images are enlarged to show merged and individual fluorescence channels. (**B**, **D** and **F**) Normalized line-scan fluorescence profiles along the dashed lines in the enlarged merged images in (**A**), (**C**) and (**E**), respectively. Scale bars, 15 μm.

**Fig. S6.**
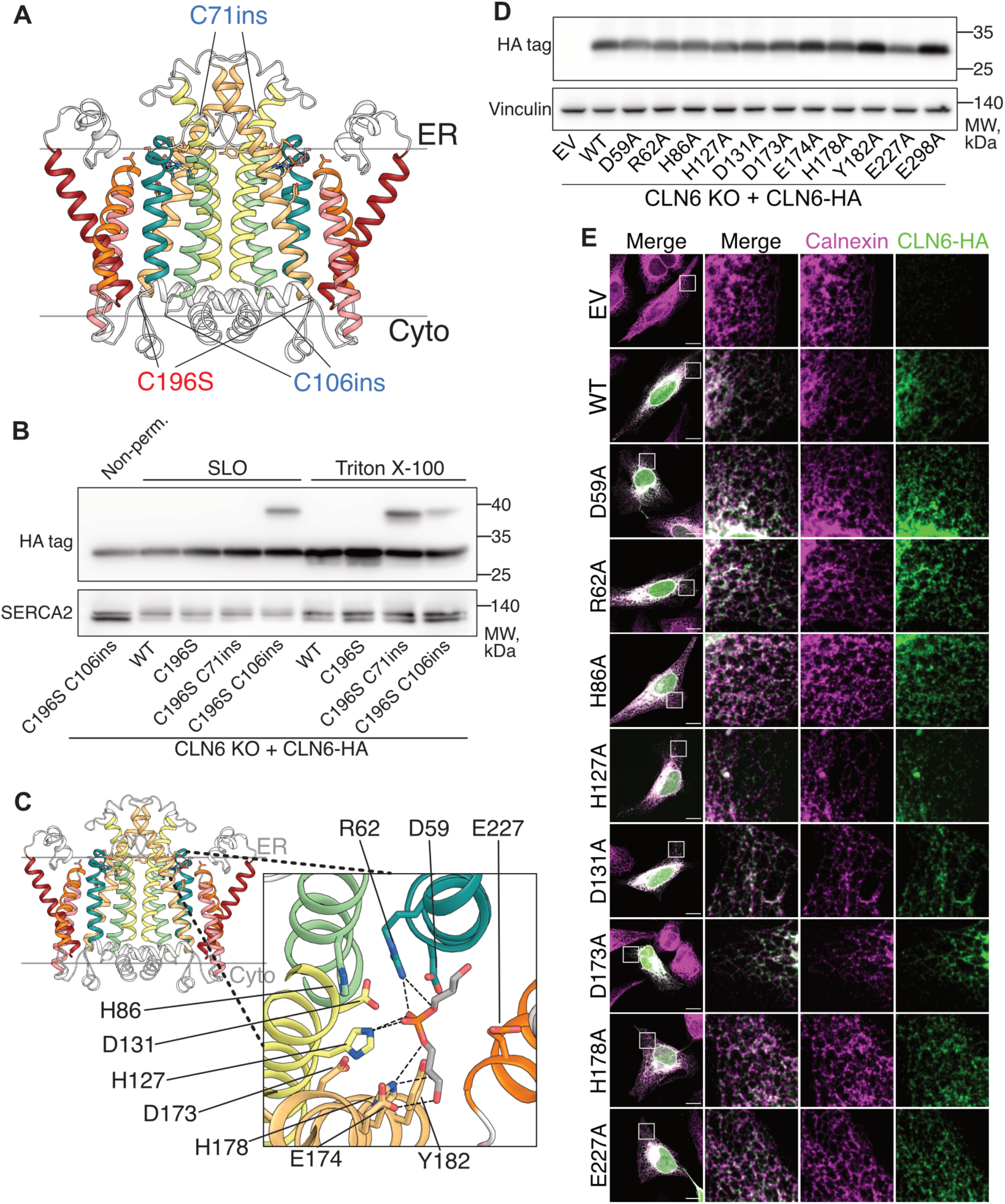
CLN6 topology mapping and alanine scan of the putative substrate-binding site. (**A**) AlphaFold 3 model of the CLN6 homodimer showing native Cys196, which was replaced with serine to reduce background maleimide labelling, and the cysteine insertion sites at positions 71 and 106, predicted to face the ER lumen and cytosol, respectively. (**B**) Topology mapping by accessibility to 5-kDa PEGylated maleimide. CLN6 KO cells expressing the indicated CLN6-HA constructs were labeled under non-permeabilized conditions, after selective plasma-membrane permeabilization with streptolysin O (SLO), or after full permeabilization with 0.1% Triton X-100. Cysteine-insertion constructs were generated on the C196S background. Anti-HA immunoblotting detects the mobility shift resulting from PEGylation; SERCA2 is shown as an ER membrane protein control. (**C**) CLN6 structural model with an enlarged view of docked *R,S*-GPG and the surrounding residues selected for alanine substitution. Dashed lines indicate putative protein–substrate contacts. (**D**) Anti-HA immunoblot showing expression of WT CLN6-HA and the indicated alanine mutants in CLN6 KO HeLa cells, with empty vector (EV) as a negative control and vinculin as a loading control. (**E**) Representative confocal immunofluorescence images of CLN6 KO HeLa cells transfected with EV, WT CLN6-HA or the indicated HA-tagged alanine mutants, showing calnexin (magenta), CLN6-HA (green) and merged signals. Regions of interest (15 x 15 μm) are enlarged in the adjacent columns. Scale bars, 15 μm.

**Fig. S7.**
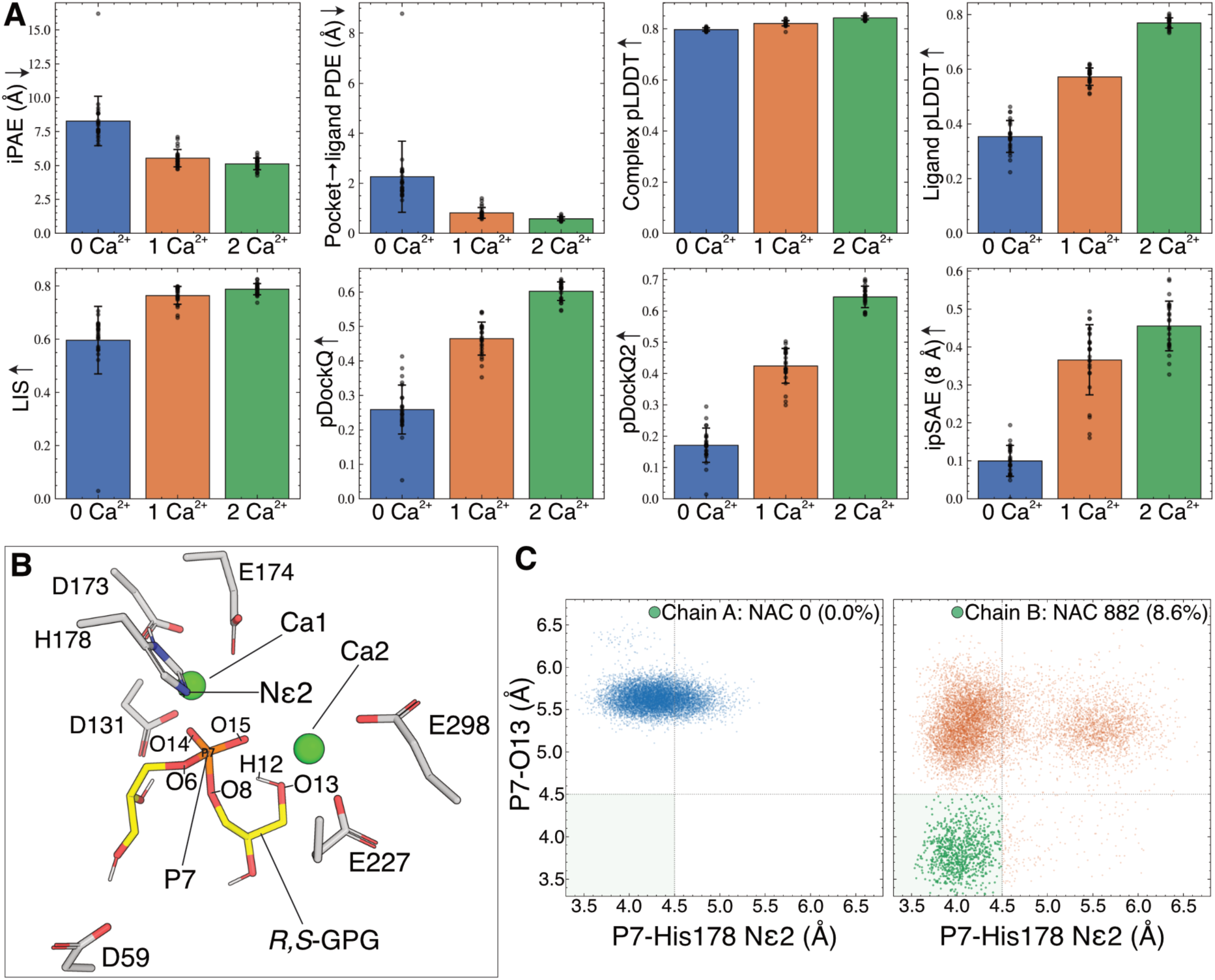
Modelling of Ca^2+^ binding and reaction-competent substrate orientation in CLN6. (**A**) Confidence metrics for CLN6–*R,S*-GPG complexes modelled with zero, one or two Ca^2+^ ions per active site. Bars and error bars indicate means ± SD across 25 models per condition; dots represent individual models. The top row shows Boltz-derived metrics: interface predicted aligned error (iPAE), pocket-to-ligand predicted distance error (PDE), and predicted local distance difference test (pLDDT) scores for the whole complex and ligand alone. Pocket residues were defined as those with a heavy atom within 8 Å of GPG. The bottom row shows additional interface-confidence scores: local interaction score (LIS), pDockQ, pDockQ2 and ipSAE, with an 8 Å PAE cutoff used for ipSAE. Arrows indicate the direction of increasing confidence. (**B**) Active site of the selected two-Ca²⁺ model, showing *R,S*-GPG with yellow carbon atoms, Ca^2+^ ions in green and CLN6 residues with grey carbon atoms. Substrate atoms and His178 Nε2 are labeled to identify the atoms used in the reaction-coordinate analyses. (**C**) Joint distributions of the distances from the substrate phosphorus (P7) to His178 Nε2 and to the distal glycerol oxygen (O13) during 500 ns of unrestrained molecular dynamics. The two protomers contained opposite substrate orientations, with the *R*-glycerol facing Asp59 in chain A and the *S*-glycerol facing Asp59 in chain B. Each point represents one analyzed trajectory frame. The shaded region defines near-attack conformations (NACs), in which both distances are ≤4.5 Å. Chain A sampled no NACs, whereas chain B sampled 882 of 10,200 frames (8.6%), highlighted in green.

**Fig. S8.**
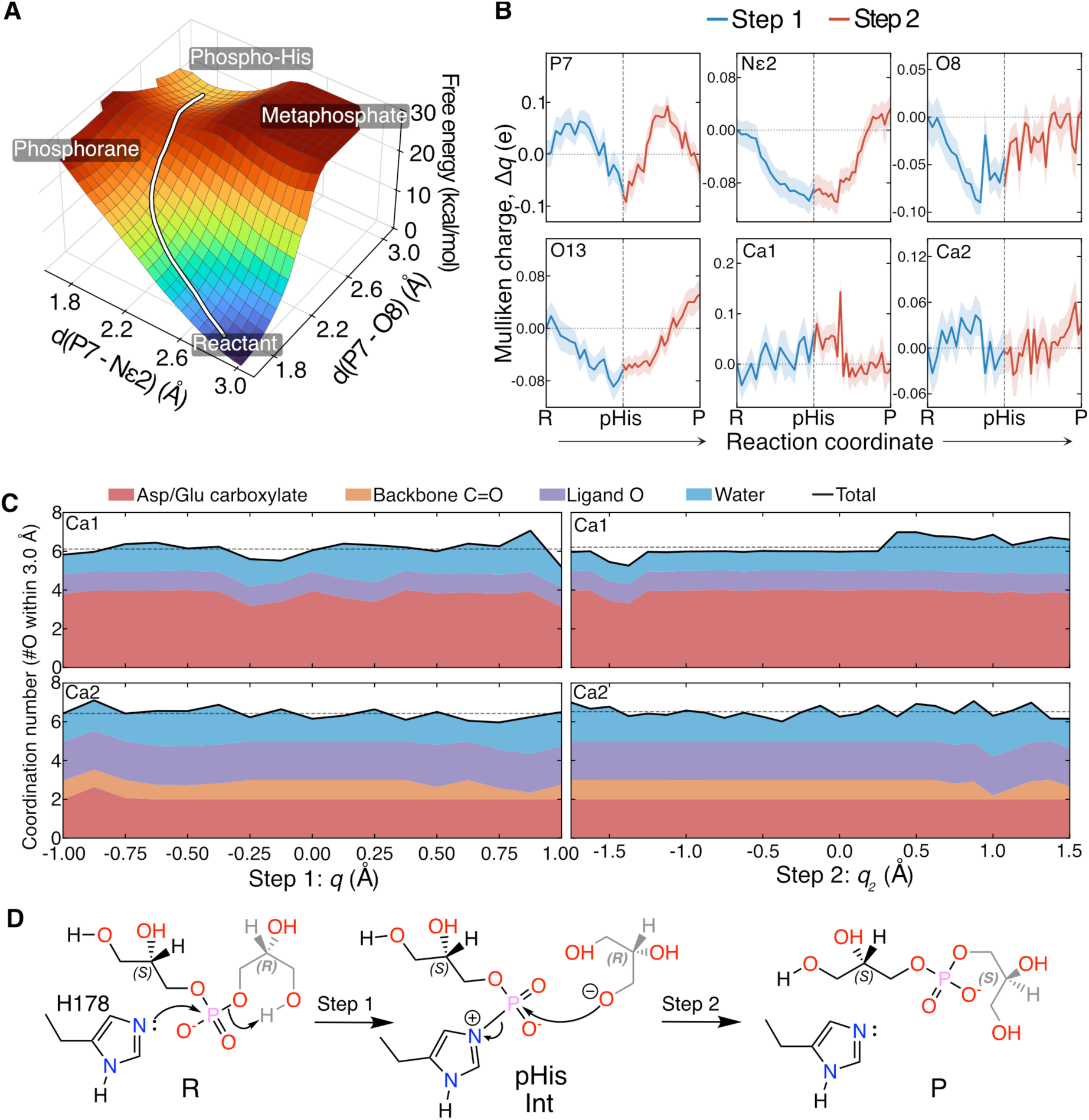
Quantum mechanics/molecular mechanics (QM/MM) analysis of the proposed His178-mediated phosphoryl transfer mechanism. (**A**) Three-dimensional representation of the free-energy surface for the first phosphoryl transfer, plotted against the forming P7–His178 Nε2 and breaking P7– O8 bond distances. The white curve indicates the proposed pathway from *R,S*-GPG to the phospho-His178 intermediate. The surface is consistent with a concerted, associative S_N_2-type transfer without a resolved phosphorane or metaphosphate intermediate. (**B**) Changes in Mulliken atomic charges (Δq, in elementary charge units, e) along the first (blue) and second (red) reaction steps for the substrate phosphorus (P7), His178 Nε2, the leaving and incoming oxygen atoms (O8 and O13), and the two structural Ca^2+^ ions. The vertical dashed line marks the phospho-His178 intermediate. R, reactant; pHis, phosphohistidine intermediate; P, product. Atom labels correspond to those in fig. S7B. (**C**) First-shell coordination of Ca1 and Ca2 during the first (left) and second (right) reaction steps. Coordination numbers were calculated from oxygen atoms within 3.0 Å of each Ca^2+^ ion and averaged within each umbrella-sampling window. Stacked areas distinguish contributions from Asp/Glu carboxylates, backbone carbonyls, GPG oxygens and water. Solid black lines show total coordination numbers; horizontal dashed lines indicate their means across each reaction step. (**D**) Proposed two-step mechanism. His178 attacks the substrate phosphorus, cleaving the original phosphoester bond and forming a covalent phosphohistidine intermediate. Attack by the alternative terminal oxygen of the released glycerol then displaces His178, yielding *S,S*-GPG.

**Fig. S9.**
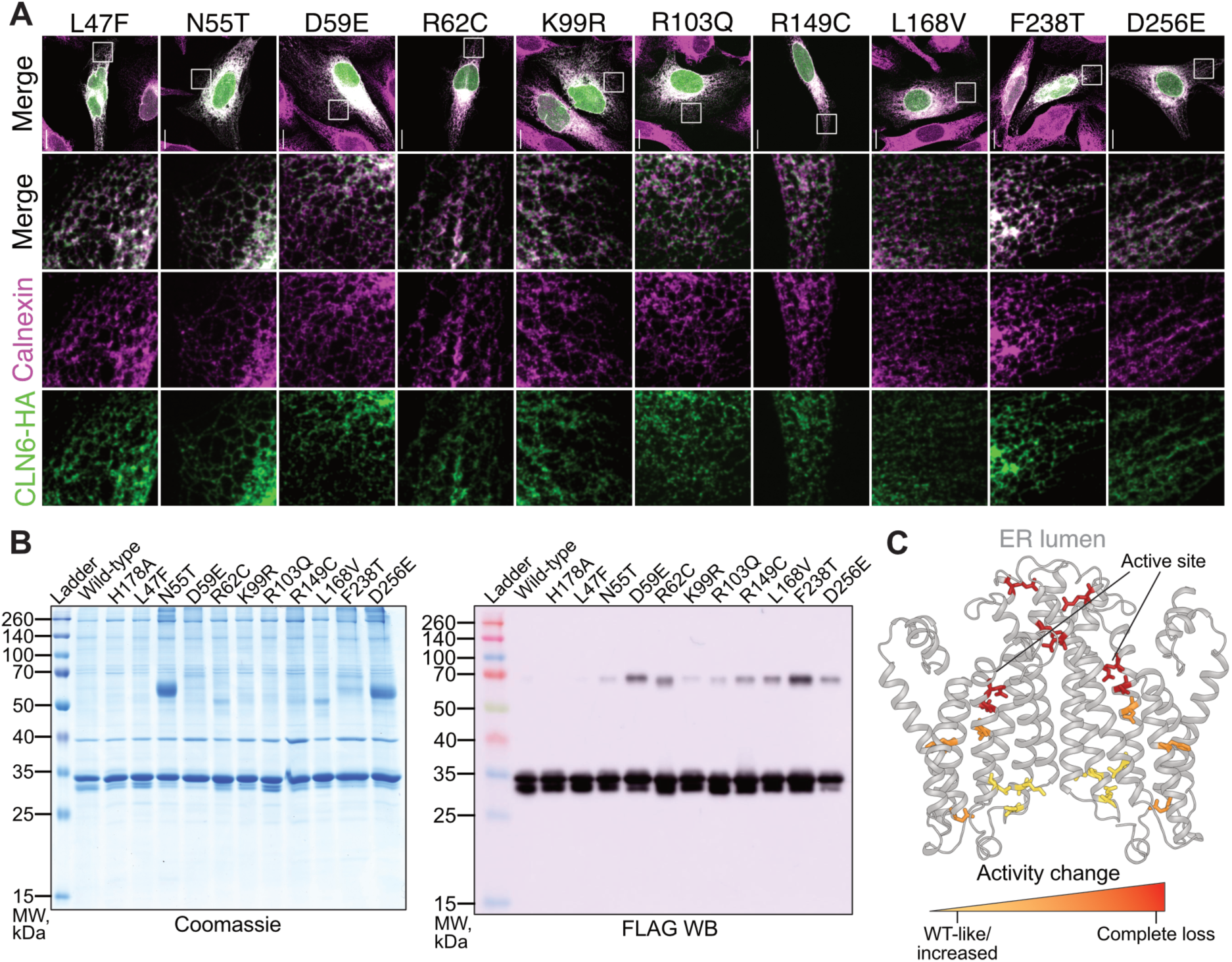
Molecular characterization of disease-associated CLN6 variants. (**A**) Representative confocal immunofluorescence images of CLN6 KO HeLa cells expressing the indicated disease-associated CLN6-HA variants. Calnexin (magenta), HA (green) and merged signals are shown. Regions of interest (15 x 15 μm) in the overview images are enlarged below. Scale bars, 15 μm. (**B**) Representative Coomassie-stained gel and anti-FLAG immunoblot of purified WT CLN6-FLAG-STREP, the catalytically inactive H178A control and the indicated disease-associated variants. Each protein lane contained 7 μg of purified protein. (**C**) AlphaFold 3 model of the CLN6 homodimer showing the positions of disease-associated substitutions, colored according to their effects on in vitro stereoconversion activity (Fig. 3E). Yellow indicates WT-like or increased activity, whereas orange to red indicates progressively greater loss of activity. The active sites and ER lumen-facing surface are indicated.

**Fig. S10.**
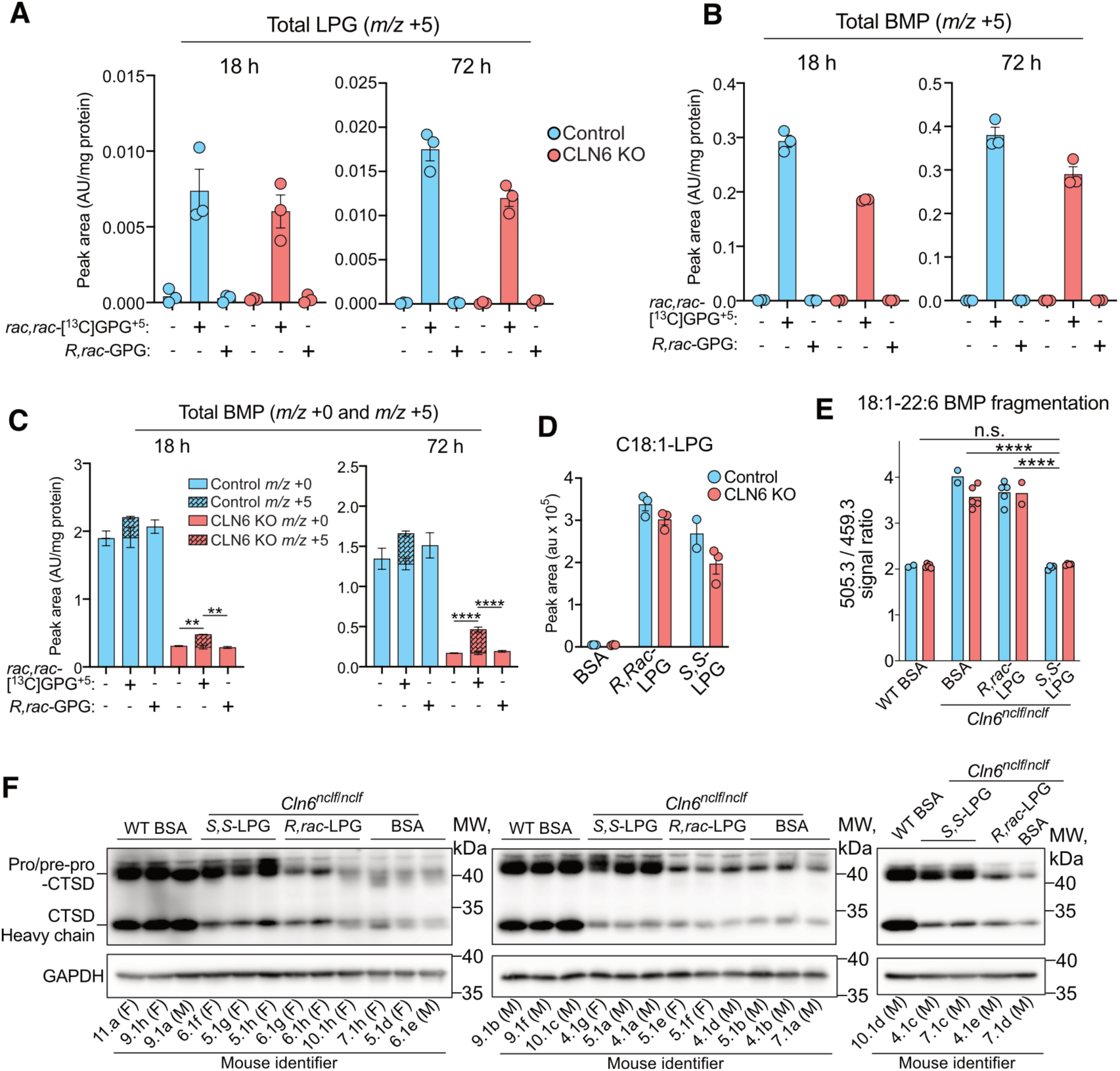
Characterization of BMP precursor rescue in cells and mice. (**A** and **B**) Abundance of ^13^C_5_-labeled LPG (**A**) and BMP (**B**) in control and CLN6 KO HeLa cells cultured without supplementation or with 250 μM *rac,rac*^13^C_5_-GPG or unlabeled *R,rac*-GPG for 18 or 72 hours. (**C**) Total BMP abundance in the same cells, partitioned into unlabeled (solid) and ^13^C_5_-labeled (hatched) fractions. Signals in (**A**) to (**C**) are normalized to cellular protein content. (**D**) C18:1-LPG abundance in control and CLN6 KO HeLa cells treated for 16 hours with BSA, 10 μM *R,rac*-C18:1-LPG or 10 μM *S,S*-C18:1-LPG. Cell experiments comprised three biological replicates per condition. (**E**) Fragmentation of asymmetric 18:1-22:6 BMP in mouse liver 16 hours after intravenous administration of BSA or 2 μmol of BSA-conjugated *R,rac*-C18:1-LPG or *S,S*-C18:1-LPG. The sodium-adducted precursor ([M+Na]^+^; *m/z* 843.6) yielded sodium-containing product ions at *m/z* 505.3 and 459.3, corresponding to the C22:6- and C18:1-containing fragments, respectively. Their signal ratio (505.3/459.3) is shown. Blue and red symbols indicate female and male mice, respectively. Measurements in (**A**) to (**E**) were performed by targeted LC-MS/MS. (**F**) Liver immunoblots of CTSD and the GAPDH loading control from the same mouse experiment, with precursor and processed heavy-chain forms of CTSD indicated. Mouse identifiers and sex are shown below each lane. The middle immunoblot is also shown in Fig. 4I. Groups in (**E**) and (**F**) comprised BSA-treated wild-type mice (n = 7; 2 female/5 male), BSA-treated *Cln6^nclf/nclf^*mice (n = 7; 2 female/5 male), and *Cln6^nclf/nclf^* mice treated with *R,rac*-LPG (n = 7; 5 female/2 male) or *S,S*-LPG (n = 8; 4 female/4 male). Bars and error bars indicate means ± SEM; points represent individual biological replicates or mice. For (**C**), \*\**P* < 0.01 and ****P < 0.0001 by two-way ANOVA with Šídák’s multiple-comparisons test. For (**E**), *P* < 0.0001 (****) by two-way ANOVA with treatment group and sex as factors, followed by Tukey’s multiple-comparisons test between treatment groups; n.s., not significant.

**Fig. S11.**
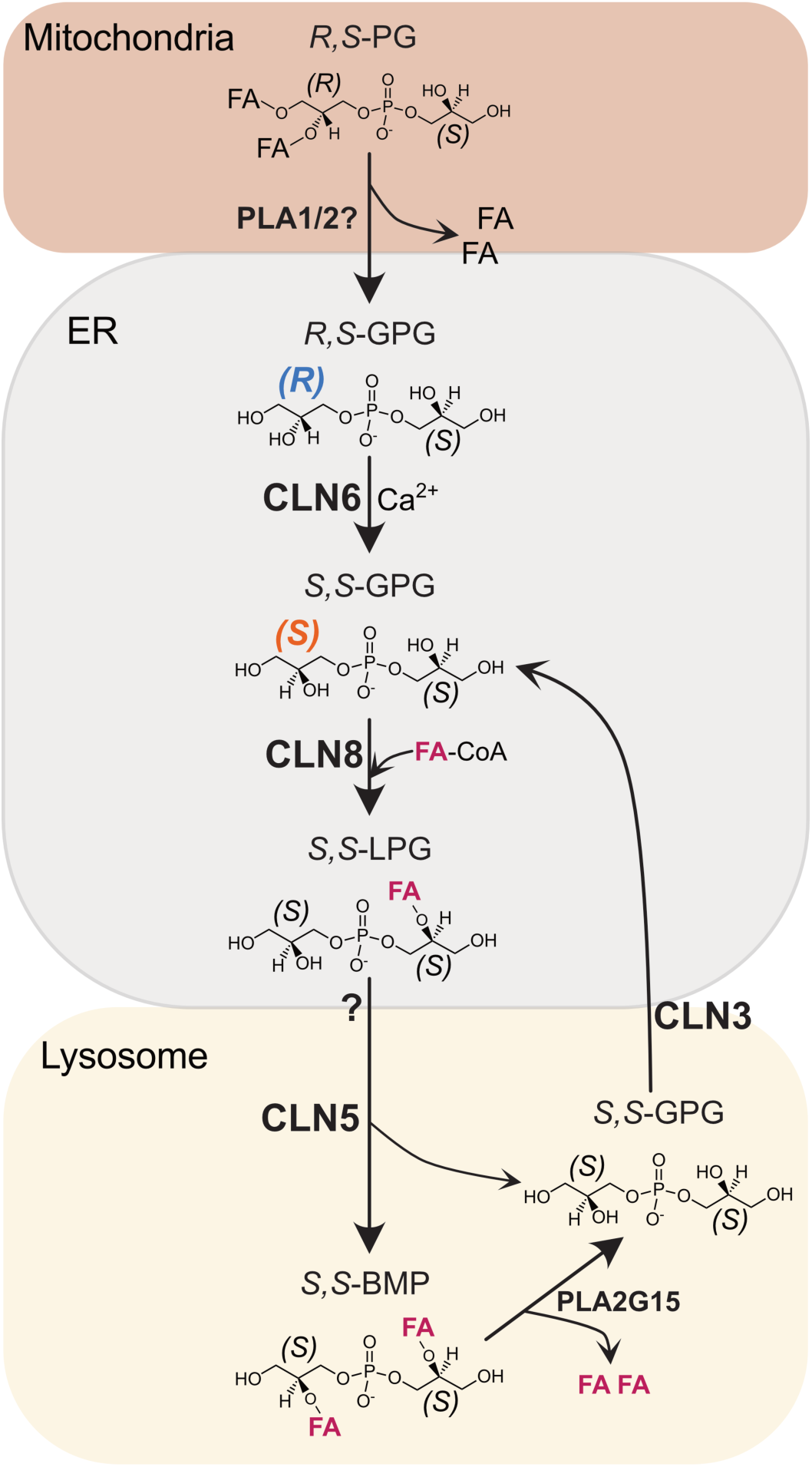
Proposed pathway of BMP synthesis and recycling. *R,S*-phosphatidylglycerol (PG) is deacylated to *R,S*-glycerophosphoglycerol (GPG) by unidentified phospholipases. CLN6 catalyzes the Ca^2+^-dependent conversion of *R,S*-GPG to *S,S*-GPG, establishing the stereochemistry of BMP. CLN8 then uses acyl-CoA to produce *S,S*-lysophosphatidylglycerol (LPG). Following delivery to lysosomes, CLN5 transfers an acyl chain between two *S,S*-LPG molecules, generating *S,S*-BMP and *S,S*-GPG. BMP catabolism by PLA2G15 also releases *S,S*-GPG. CLN3-dependent lysosomal export (*35*) allows *S,S*-GPG to be reused by CLN8, maintaining its stereochemistry through successive rounds of synthesis and turnover. Question marks indicate the unidentified phospholipases responsible for PG deacylation and the unresolved mechanism of LPG delivery to lysosomes. Colored backgrounds indicate the organelles associated with the pathway steps. FA, fatty acid.

